# Cryo-EM structure of the complete *E. coli* DNA Gyrase nucleoprotein complex

**DOI:** 10.1101/696609

**Authors:** Arnaud Vanden Broeck, Julio Ortiz, Valérie Lamour

## Abstract

DNA Gyrase is an essential enzyme involved in the homeostatic control of DNA supercoiling and the target of successful antibacterial compounds. Despite extensive studies, the detailed architecture of DNA Gyrase from the model genetic organism *E. coli*, is still missing, impeding structure-function analysis of *E. coli-*specific catalytic regulation and limiting the study of conformational intermediates of this highly flexible macromolecule. Herein, we determined the complete molecular structure of the *E. coli* DNA Gyrase bound to a 180 bp DNA and the antibiotic Gepotidacin, using phase-plate single-particle cryo-electron microscopy. Our data unveil with unprecedented details the structural and spatial organization of the functional domains, their connections and the position of the conserved GyrA-box motif. The deconvolution of closed and pre-opening states of the DNA-binding domain provides a better understanding of the allosteric movements of the enzyme complex. In this region, the local atomic resolution reaching up to 3.0 Å enables the identification of the antibiotic density in the DNA complex. Altogether, this study paves the way for the cryo-EM determination of gyrase complexes with antibiotics and opens perspectives for targeting conformational intermediates. The type 2A DNA topoisomerases (Top2) are nanomachines that control DNA topology during multiple cellular processes such as replication, transcription and cell division ^1-4^. These enzymes catalyze the transport of a DNA duplex through a double strand break to perform DNA relaxation, decatenation and unknotting. DNA Gyrase plays a vital role in the compaction of the bacterial genome and is the sole type 2 topoisomerase able to introduce negative supercoils into DNA, a reaction coupled to ATP hydrolysis ^5^.

---

DNA Gyrase A and B subunits assemble into a A_2_B_2_ hetero-tetramer of approximatively 370 kDa forming 3 molecular interfaces called N-gate, DNA-gate and C-gate allowing DNA binding and strand passage ^6,7^. The flexibility of this large enzyme constitutes a challenge for the structural study of the multiple conformations it adopts during the catalytic cycle. Until recently, only the structures of isolated domains with drugs were known, providing partial information on these modular enzymes and not accounting for the allosteric connections between the catalytic domains ^8-11^. The architecture of a full-length DNA Gyrase from *T. thermophilus* in complex with a 155 bp dsDNA and ciprofloxacin was first solved using cryo-electron microscopy (cryo-EM) at a resolution of 18 Å ^12^. This first model provided a rationale for probing mechanistic questions based on biochemical, single molecule and FRET techniques ^13-16^, but the low resolution model did not allow deconvolution of discreet conformations of the full-length enzyme, leaving many mechanistic questions unresolved. Furthermore, the atomic details of the structure and of the drug binding site in the context of the overall conformation of DNA gyrase were not available at this resolution.

In addition, the information derived from the thermophilic homolog does not account for all the mechanistic and structural specificities of the *E. coli* DNA gyrase, the genetic model for which functional data has been accumulated over the past decades. In particular, the *E. coli* DNA Gyrase GyrB subunit possesses a domain insertion of 170 amino acids, not found in the *T. thermophilus* enzyme, that has been shown to help coordinate communication between the different functional domains ^17^. Deletion of this domain greatly reduces the ability of *E. coli* DNA Gyrase to bind DNA and decreases its ATP hydrolysis and DNA negative supercoiling activity ^17^. DNA Gyrase is a prime target for catalytic inhibitors such as aminocoumarins, or ‘poison’ inhibitors of the cleavage complex, such as quinolones ^18-20^. More recently, novel bacterial topoisomerase inhibitors (NBTIs), have been developed that also target the DNA cleavage activity of Gyrase, but with a mechanism and target site different from the quinolones ^21^. These molecules and in particular Gepotidacin, represent a promising alternative and are currently in clinical trial for the treatment of bacterial infections ^22-24^. However, the use of any antibiotics may lead to the appearance of mutations and the development of resistant strains ^25^. Our capacity to understand the molecular determinants of drug resistance depends in part on the availability of complete 3D architecture of DNA gyrase nucleoprotein complexes with drugs.

In this study, we reveal the complete molecular structure of the *E. coli* DNA Gyrase bound to dsDNA and to the NBTI molecule Gepotidacin using cryo-EM. The structure of the entire complex was solved at 6.6 Å resolution and two conformations of the DNA-binding domain in closed and pre-opening states were solved at 4.0 Å and 4.6 Å resolution, respectively. These subnanometer resolution structures reveal in unprecedented detail the structural and spatial organization of the different functional domains of *E. coli* gyrase and of their connections. In particular, they elucidate the position of the conserved GyrA box motif responsible for DNA wrapping and allow the unambiguous identification of the Gepotidacin molecule in the cryo-EM map. The deconvolution of distinct conformations of the DNA-binding domains provides a better understanding of the allosteric movements that the enzyme conducts at the early steps of G-segment opening, stabilized by the Gepotidacin molecule. Altogether this work paves the way for the structure determination of gyrase complexes with additional antibiotics using cryo-EM and the in-depth analysis of its allosteric regulation.

## Results

### Formation of the DNA-bound Gyrase complex and cryo-EM 3D reconstruction

The *E. coli* GyrB and GyrA subunits were overexpressed and purified separately before assembly in stoichiometric amounts and incubation with a double-nicked 180 bp DNA (Supplementary Table 1). The holoenzyme was further purified by size-exclusion chromatography resulting in a single and homogenous complex (Fig. 1a and Supplementary Figure 1a). DNA supercoiling and ATP hydrolysis assays showed that the A_2_B_2_ complex is fully active (Supplementary Figure 1b). Finally, the Gepotidacin molecule was added to the complex which was further stabilized using ADPNP, the non-hydrolysable analog of ATP.

**Fig 1.**
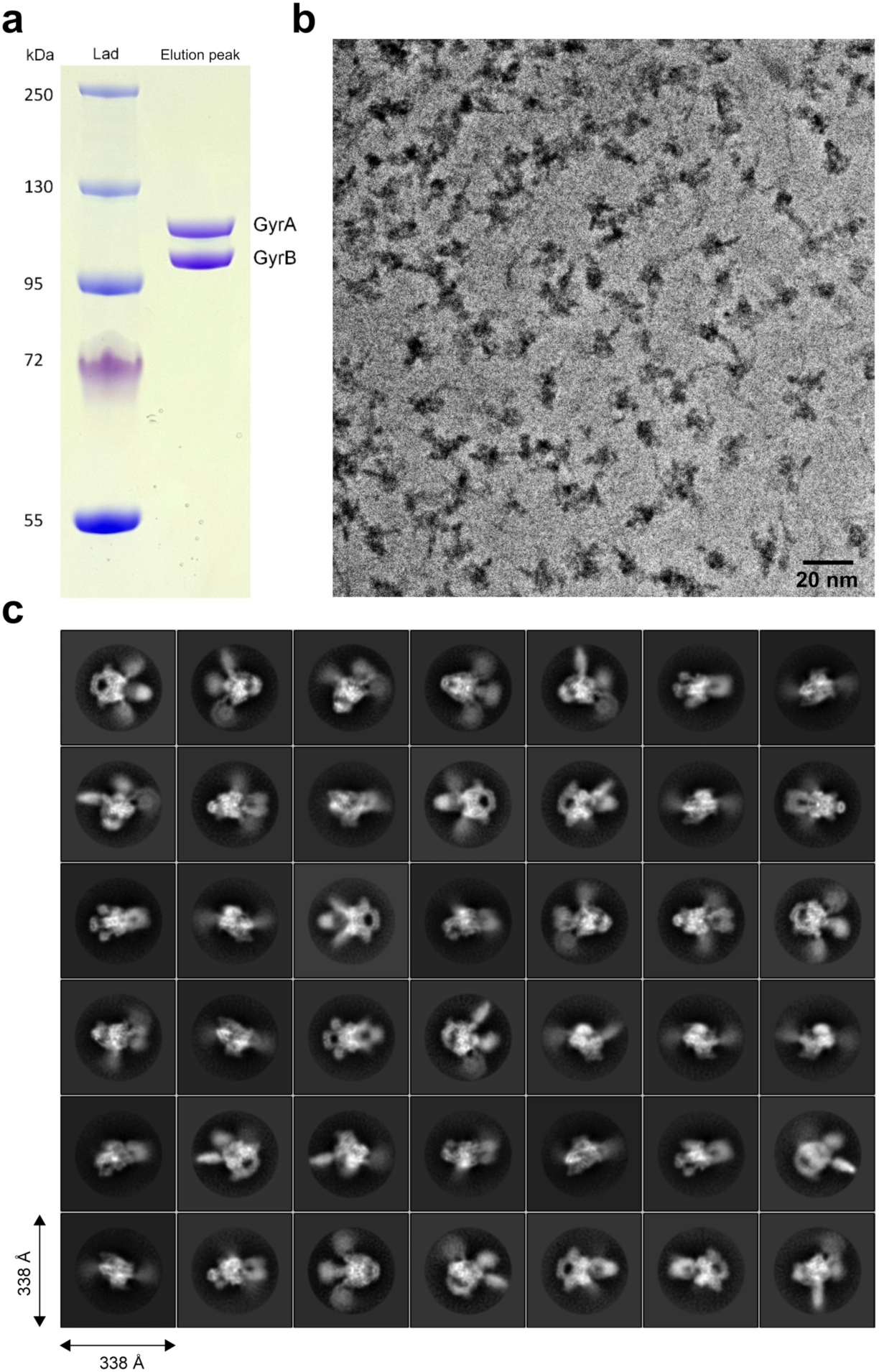
Sample preparation and image acquisition. **a**. SDS-PAGE analysis of the reconstituted GyrA_2_B_2_ after size exclusion chromatography. **b**. A typical cryo-EM micrograph collected with a Gatan K2 Summit camera on a FEI Titan Krios microscope operated at 300?kV with Volta Phase Plate. **c**. Selection of 2D classes from reference-free 2D classification.

Images of the DNA Gyrase complex were recorded using a Titan Krios with a Volta Phase-Plate at a 500 nm defocus target, allowing a significant increase in contrast of the images and optimization of the image acquisition rate ^26,27^ (Fig. 1b). After gain correction, frame alignment and manual inspection, 8,701 micrographs were selected out of 11,833 initial movie frames coming from 4 datasets (Supplementary Table 2). Several rounds of 2D classification in RELION2 ^28,29^ yielded class averages displaying high-resolution features such as the DNA wrapped around the β-pinwheels and defined alpha helices (Fig. 1c). An *ab-initio* 3D model was calculated with a final stack of 191,456 particles using cryoSPARC ^30^ (Supplementary Figure 2). Multiple classification and refinement steps were then performed in RELION2 using global and local approaches to deconvolute different structures of the DNA-bound DNA Gyrase (Supplementary Figure 3-4). The overall DNA-bound DNA Gyrase complex was solved at 6.6 Å using 94,633 particles. The estimation of the local resolution showed a broad spectrum from 5 Å on the DNA-binding core to 8 Å on the ATPase domains and β-pinwheels (Supplementary Figure 4c).

**Fig 2.**
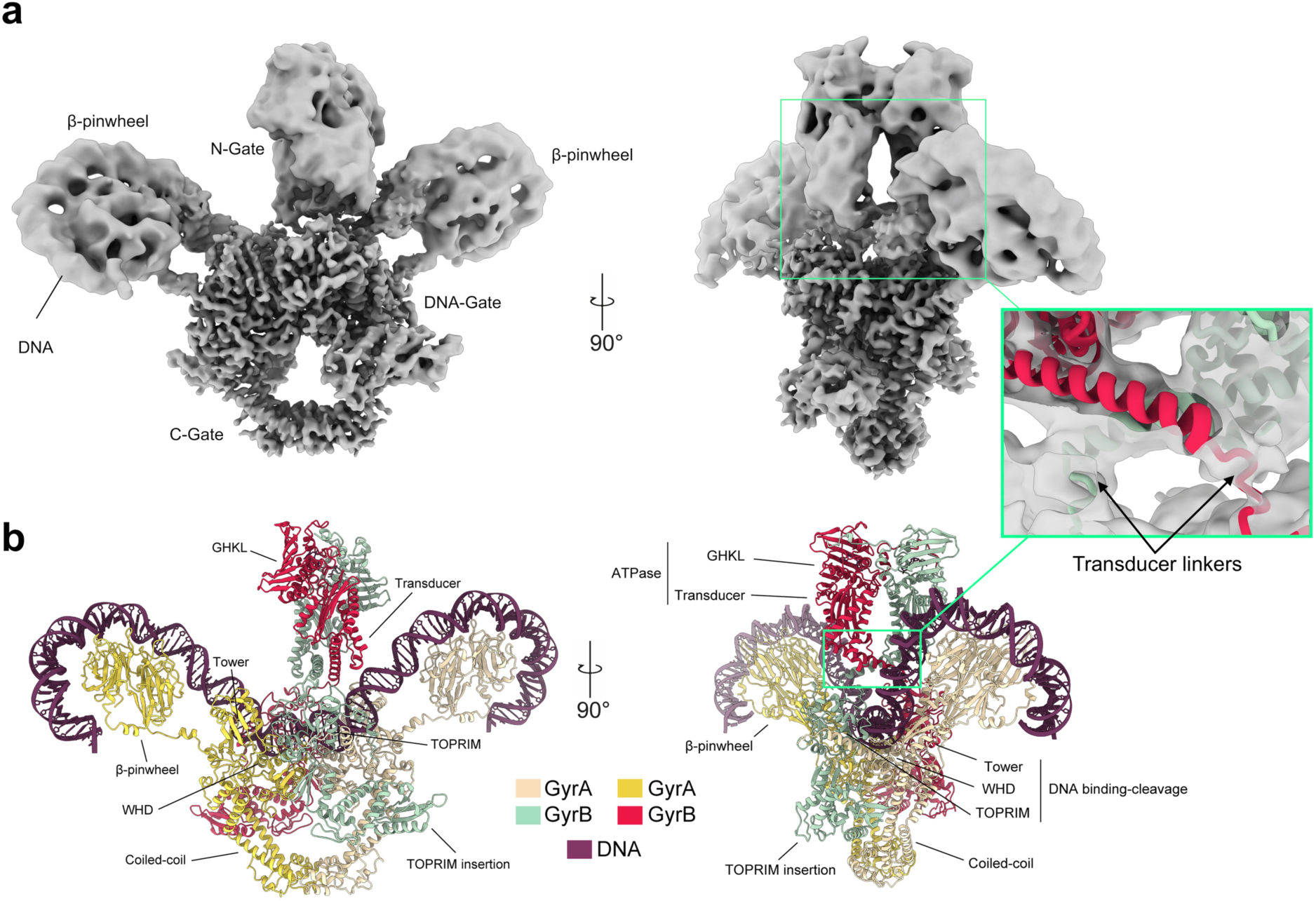
Cryo-EM 3D reconstruction and molecular model of the DNA Gyrase complex. **a**. Composite cryo-EM map of the full complex of DNA Gyrase with an inset showing the slabbed density of the transducer helices connecting the ATPase domain (N-gate) to the DNA cleavage-domain (DNA-gate). **b**. Molecular structure of the heterotetrameric complex including the GyrA/GyrB subunits and 130 bp of DNA that were built and refined in the cryo-EM maps (see also Supplementary Figures 3 and 5).

**Fig 3.**
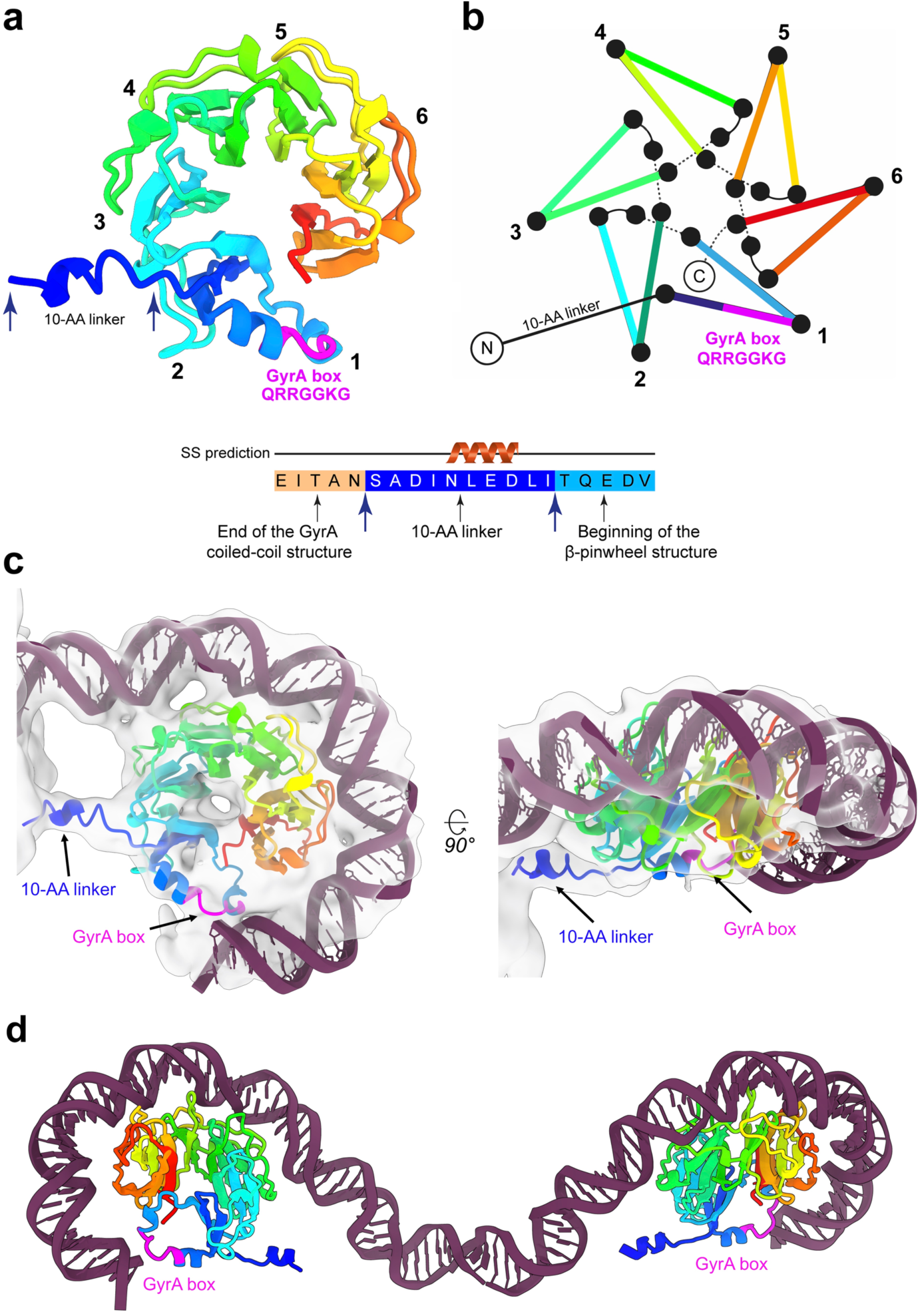
DNA Wrapping around the GyrA CTD β-pinwheel and GyrA box structure. **a**. Cartoon representation of the molecular structure of the GyrA β-pinwheel completed in this work, (rainbow colored from the N-terminal end in blue, to the C-terminal end in red. The GyrA box (QRRGGKG) is colored in magenta. The β-pinwheel blades are numbered from 1 to 6. **b**. Schematic representation of the β-pinwheel with the same color code and numbering. **c**. Two orthogonal views of the β-pinwheel wrapped with DNA (in purple) in the 6.3 Å cryo-EM map (grey surface). The identification of the 10-AA linker in the map allowed positioning of the β-pinwheel structure unambiguously. The linker connecting the β-pinwheel with the GyrA coiled-coil domain attaches to the lower part of the β-pinwheel. The first contact between DNA and the β-pinwheel occurs at blade 3, wrapping around the β-pinwheel by contacting blade 4, 5, 6 and exiting the β-pinwheel through contact with blade 1. This leads to the positioning of the GyrA box (in magenta) at the exit of the DNA path around the β-pinwheel, in close contact with DNA. **d**. Overall view of the 130bp DNA duplex wrapping around the two β-pinwheels of the DNA Gyrase. For clarity, the rest of the structure is not shown. GyrA box motifs of each β-pinwheel are situated at the exit of the DNA path around the β-pinwheel and act as clamps stabilizing the DNA curvature.

**Fig 4.**
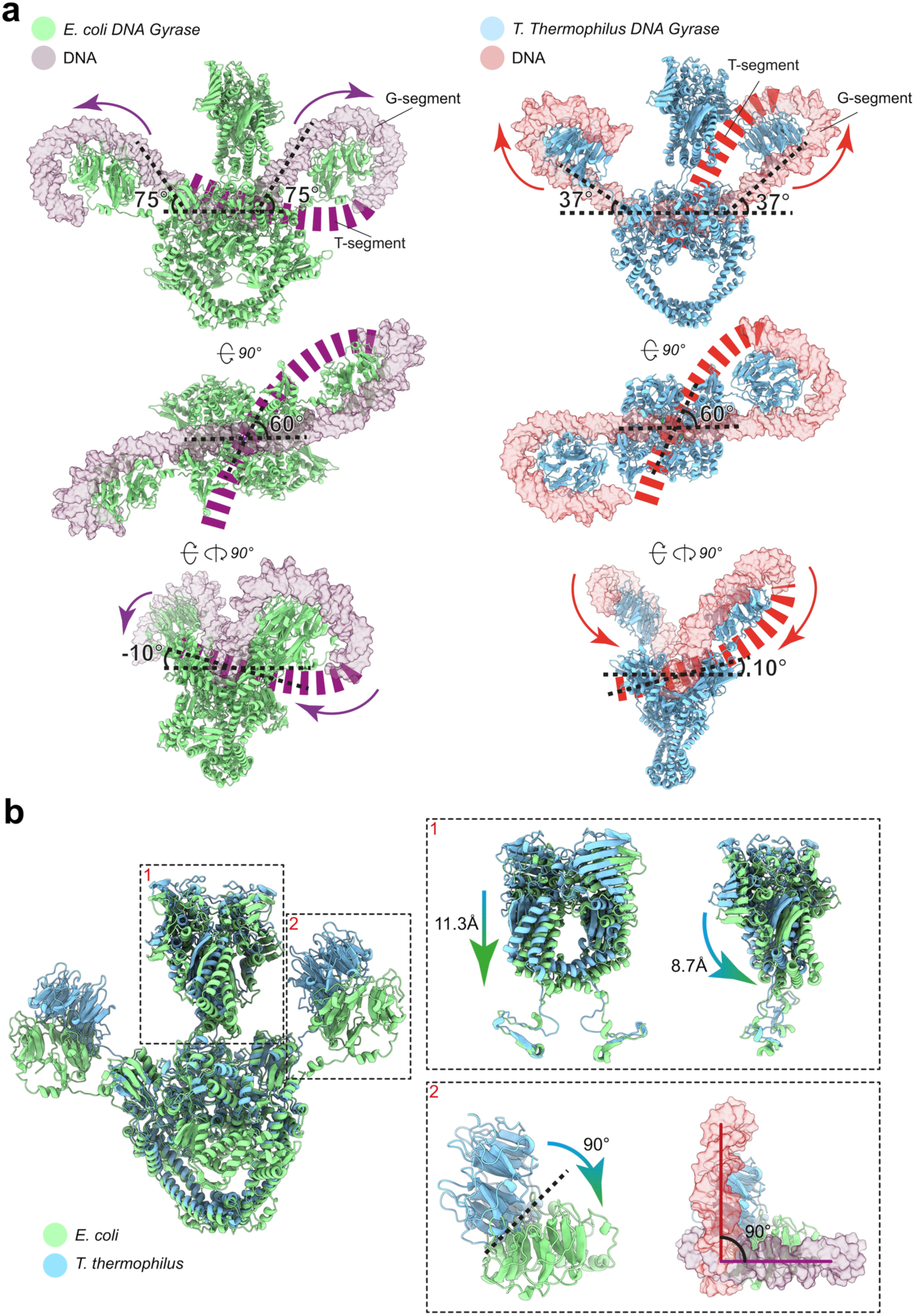
Structural comparison of the *E. coli* and *T. thermophilus* DNA Gyrase bound to DNA. **a**. Three orthogonal views of the *E. coli* and *T. thermophilus* ^12^DNA Gyrase structures bound to DNA. In the top and side views, the ATPase domain has been omitted for clarity. The spatial arrangement of the β-pinwheels induces an overall ∼150° bending of the DNA for *E. coli* DNA Gyrase and ∼75° for *T. thermophilus* DNA Gyrase. In both structures, the 130 bp DNA duplex (purple and red) is chirally wrapped around the β-pinwheel in an orientation that is consistent with the formation of a 60° angle positive crossover when the T-segment path is extrapolated in the DNA-gate groove formed by the TOPRIM-WHD and TOWER domains. The angle of the T-segment entering the Gyrase core is different between the two structures: −10° for *E. coli* and 10° for *T. thermophilus*. **b**. Superimposition of the *E. coli* and *T. thermophilus* DNA Gyrase. The insets show the structural differences. (1) The *E. coli* ATPase domain shits towards the DNA-binding and cleavage domain by ∼11 Å compared to *T. thermophilus*. (2) The β-pinwheels are separated by a 90° rotation angle, which induces changes in the overall bending of the DNA and the angle of attack of the T-segment.

Structures of the DNA-binding and cleavage domain were solved in two different states with different positioning of the GyrB TOPRIM insertion relative to the catalytic domains. State 1 was solved at 4.0 Å using 60,548 particles and State 2 was solved at 4.6 Å using 53,655 particles (Supplementary Figure 3). Local resolution estimation shows more stable regions reaching up to 3 Å and 4 Å for State 1 and State 2 structures respectively, while the GyrB *E. coli* insertion seems more disordered (Supplementary Figure 4c). To allow better positioning and refinement of the atomic models in the EM density of the overall structure, we also solved three additional structures with better defined densities for the ATPase head and DNA-binding core (5.9 Å), the DNA-binding core without the TOPRIM insertion (4.0 Å) and the C-terminal β-pinwheels wrapped by DNA (6.3 Å) (Supplementary Figure 3-5 and Supplementary Table 2). The cryo-EM maps of individual regions at different resolutions were used to generate a composite EM map that reflects the flexibility of the complex, in particular in the ATPase and DNA-bound β-pinwheels (Fig 2a, Supplementary Figure 5 and Supplementary Movie 1).

**Fig 5.**
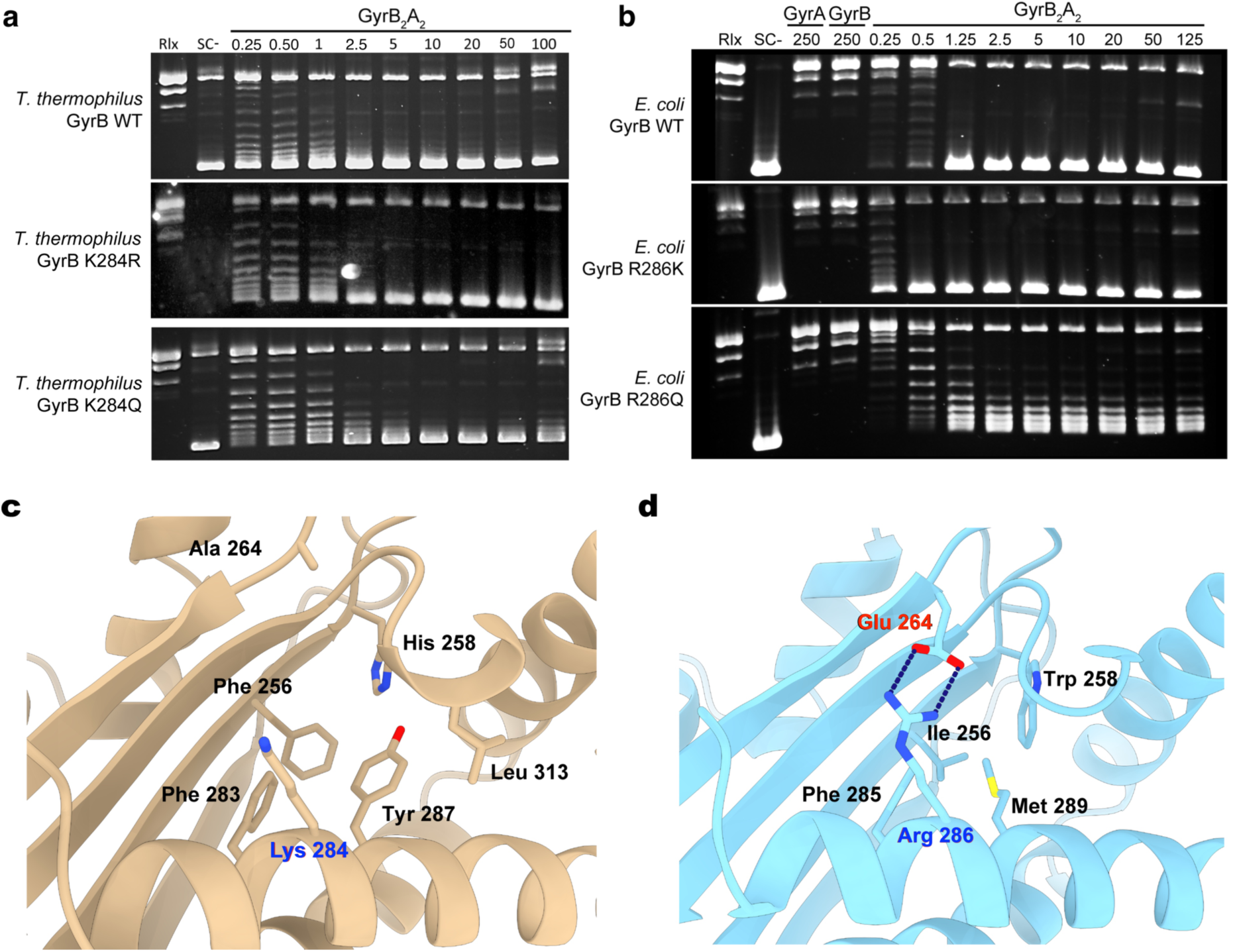
Transducer structural elements involved in allosteric regulation. **a**. DNA negative supercoiling activities of wild-type, K284R and K284Q *T. thermophilus* DNA Gyrase showing no effect of the mutations. Protein concentrations are indicated in nM holoenzyme. Negative and positive controls are shown as relaxed (Rlx) or negatively supercoiled DNA species (SC-), respectively. **b**. DNA negative supercoiling activity of wild-type, R286K and R286Q *E. coli* DNA Gyrase showing no effect of R286K but an impaired activity of the R286Q mutant. **c**. Cartoon representation of the *T. thermophilus* GyrB transducer domain. The K284 residue is not engaged in an interaction network. The transducer is mainly stabilized by a hydrophobic core (F256, F283, Y287, L313). **d**. Cartoon representation of the *E. coli* GyrB transducer domain. The transducer domain is stabilized by a salt bridge involving R286 and E264 residues anchoring the beta-sheets to the alpha helix.

### Completeness of the *E. coli* DNA Gyrase model and analysis of the DNA complex architecture

The atomic models of the GyrA and GyrB domains solved by X-ray crystallography ^10,17,31^ were combined to build and refine the complete atomic structure of the full-length *E. coli* DNA Gyrase holoenzyme in complex with the DNA duplex in the cryo-EM maps (Fig. 2b). The continuous density of the ATPase and DNA-binding domains EM map at 5.9 Å allowed us to build unambiguously the protein linkers between the C-terminal end of the transducer helices of the ATPase domain and the N-terminal end of the TOPRIM domain (Fig. 2 and Supplementary Figure 5). We could also build the linker between the C-terminal end of the GyrA coiled-coil domain and the β-pinwheel on each side (Fig. 2 and Supplementary Figure 5). The tracing of this linker, together with the quality of the density that allows to distinguish the lower from the upper face of the pinwheel disk, enabled the precise orientation of the crystal structure of the *E. coli* β-pinwheel in the EM map ^10^ (Fig. 3). Consequently, the first contact between the G-segment and the β-pinwheel occurs at blade 3, then wraps around the β-pinwheel by contacting blade 4, 5, 6 and exits the β-pinwheel through contact with blade 1 containing the GyrA-box motif (Fig. 3c-d). Other structural elements missing in crystal structures such as surface loops, *α*-helices of the DNA binding/cleavage and C-gate domains could be completed or corrected in the GyrB and GyrA subunits representing more than 5% of the total sequence (Supplementary Figure 6 and Supplementary Table 3).

**Fig 6.**
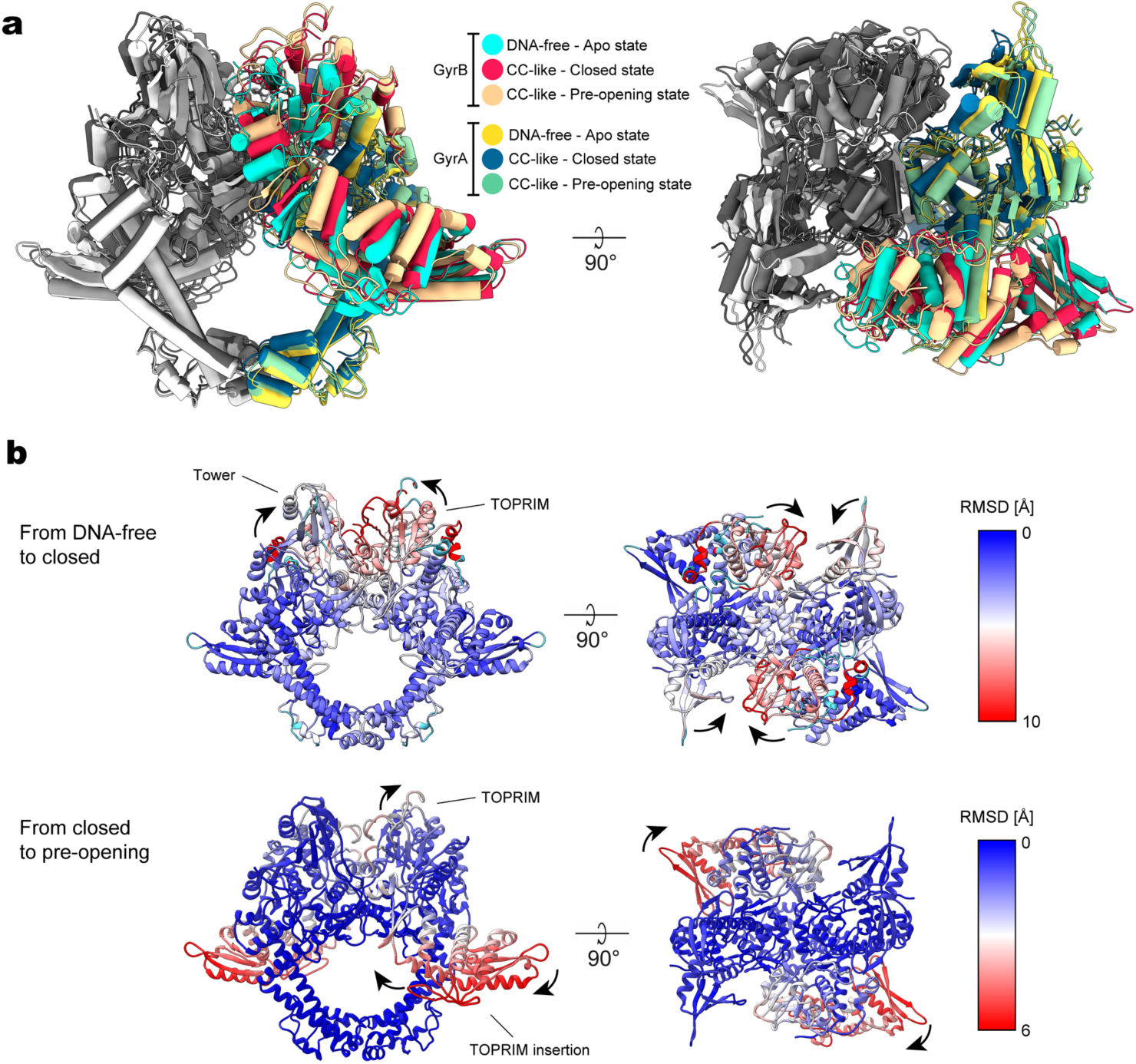
Conformational changes associated with G-segment binding, cleavage and opening. **a**. Superimposition of DNA-binding and cleavage domain structures in different states: a DNA-free form in apo state (PDB ID 3NUH), a cleavage complex in closed state (this study) and a cleavage complex in pre-opening state (this study). Only one monomer of GyrB and GyrA is colored as indicated in the legend and DNA is omitted for clarity. The superimposition shows the upward movement of the TOPRIM and Tower domains from the apo to the closed and pre-opening states. **b**. RMSD analysis of the DNA-binding and cleavage domain in *apo* state and cleavage complex in closed (state 1) and pre-opening (state 2) conformations. Upper panel: from DNA-free *apo* state to the cleavage complex state in closed conformation. The majority of the movements are performed by the TOPRIM and Tower domains upon DNA binding and cleavage. Lower panel: from the cleavage complex in closed conformation to the pre-opening conformation. The TOPRIM insertion domain pivots around the N-terminal GyrA arm preceding opening of the G-segment after cleavage.

The resulting model now shows clearly the details of the intertwined structure of the DNA Gyrase heterotetramer (Fig. 2b). The dimeric ATPase domain is sitting in an orthogonal orientation above the DNA binding domain (∼95°) as previously observed with the *T. thermophilus* DNA Gyrase ^12^ (Fig. 4a and Supplementary Figure 7). The structure is asymmetric with the ATPase domain slightly bent (∼10°) towards one of the β-pinwheels bringing it closer to a distance of 27 Å. The ATPase domain is sitting 11 Å away from the DNA binding domain (Fig. 4b). Each β-pinwheel resides on the same side as its respective GyrB subunit, with an approximate ∼45° angle from the center of the DNA-binding domain. This spatial arrangement of the β-pinwheel induces an overall ∼150° bending of the DNA, as seen in previously reported crystal structures ^21,32,33^. 130 bp of the double-stranded DNA out of 180 bp could be fitted and the corresponding base pairs were submitted in refinement in the cryo-EM map. The DNA duplex is chirally wrapped around the β-pinwheels orienting the T-segment towards the DNA-gate second groove formed by the TOPRIM-WHD and tower domains with a 60° angle.

**Fig 7.**
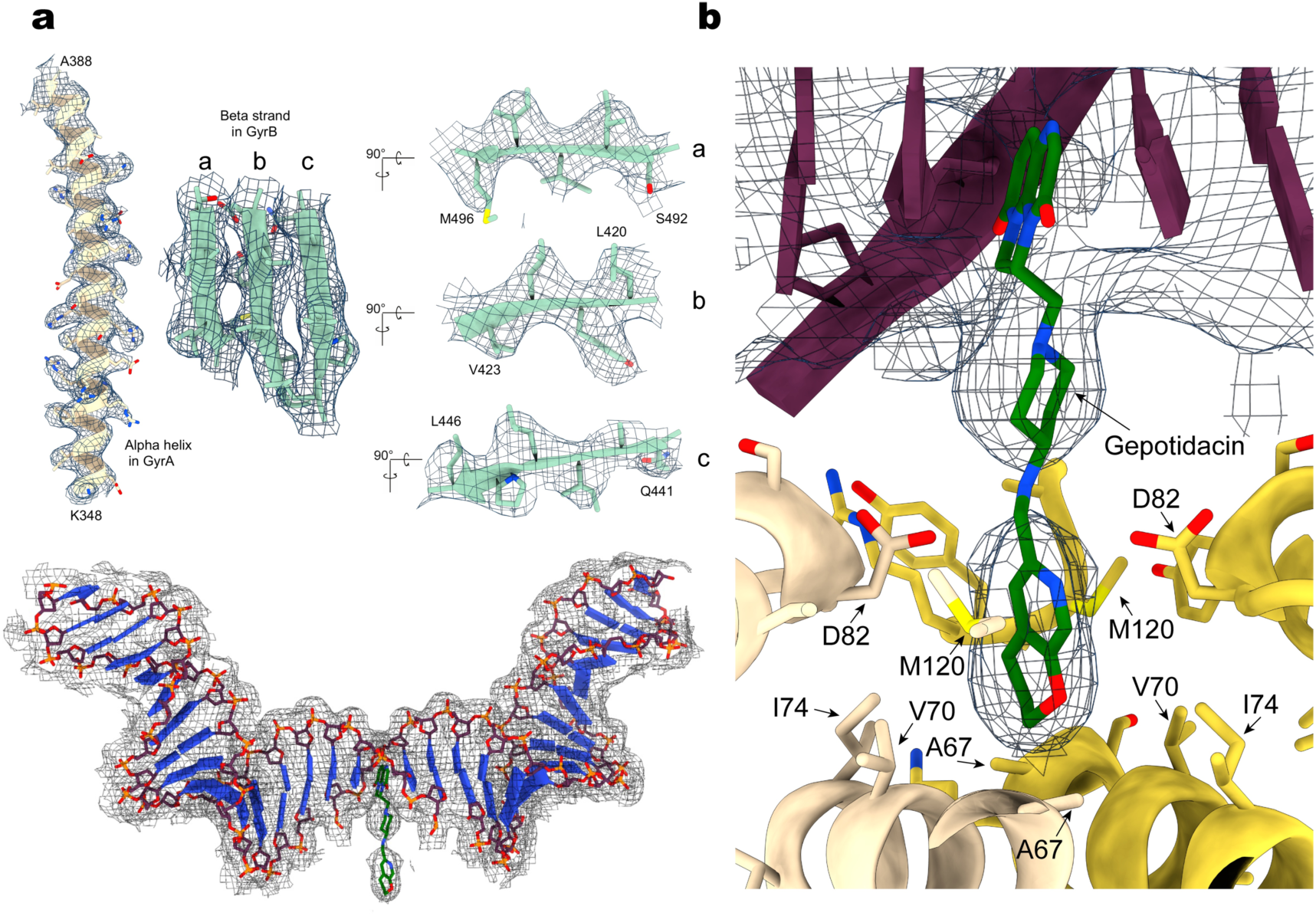
High resolution features of the DNA binding and cleavage domain bound to Gepotidacin. **a**. High resolution features of the 4 Å cryo-EM map. An alpha helix and beta-sheet with well resolved side chains are shown on the upper panel. The 36 bp DNA duplex and Gepotidacin visualized in the electronic density of the 4 Å cryo-EM map are shown in the bottom panel. **b**. Gepotidacin binding site. Electron density of the DNA and Gepotidacin are shown in blue mesh. Residues in the direct vicinity of the compound are highlighted.

### Analysis of conserved structural elements of the transducer domain involved in allosteric control

The connection between the ATPase domain and DNA binding cleavage domain is now clearly built and positioned in the gyrase model (Fig 2a). This connection is ensured by the transducer helices in the GyrB subunit which are thought to be a pivotal structure for the allosteric changes following ATP hydrolysis ^31,34^. These helices are sticking out of the transducer domain that forms a central cavity in the ATPase domain. It was suggested that residue R286 in *E. coli* (*Ec*) pointing towards the cavity of the transducer domain contributes to DNA capture and therefore is a key element of the enzyme’s allostery ^35^. This residue is replaced by a lysine in the *Thermus thermophilus* (*Tth*) enzyme and most organisms possess either an arginine or a lysine at this position (Supplementary Figure 8). To assess if this position is important for the allosteric regulation of the holoenzyme, we compared the catalytic activities of point mutations of *Ec* R286 and *Tth* K284 (Fig. 5). *Ec* R286 was mutated into a lysine to mimic the *Thermus* position and a glutamine to remove the charge. Inversely, *Tth* K284 was mutated into arginine or glutamine. The mutant proteins were recombinantly produced with no detectable denaturation. Thermal denaturation experiments on the mutant proteins using differential scanning fluorimetry showed the same behavior as the WT protein (Supplementary Figure 9a). The ATPase and DNA supercoiling activities *of the Tth* K284R and K284Q mutants remain in the same range as the WT at 37°C (Fig. 5a and Supplemental Figure 9b). The *Ec* R286K mutant is also not impaired in any of the tested activities compared to the WT suggesting either side chain is compatible with the catalytic activities in *E. coli* (Fig. 5b and Supplemental Figure 9b). In contrast, the *Ec* R286Q mutant has a 5-fold reduced negative supercoiling activity and a 6-fold reduced ATPase activity.

Analysis of the ATP binding domain structure in this area shows that *Ec* R286 is engaged in a salt bridge with E264 (Fig. 5d). This interaction anchors the β-sheet of the transducer to the *α*-helix carrying R286, providing rigidity to the transducer subdomain. In contrast, *Tth* K284 at the same position points toward the N-gate central cavity with no obvious role in the transducer stabilization. The *Thermus* transducer is instead stabilized by an hydrophobic core involving 5 aliphatic and aromatic residues (Fig. 5c). The R286-E264 salt bridge interaction in *Ec* is most likely to be disrupted by a charge inversion, explaining the loss of activity of the *Ec* R286Q mutant. As a consequence, we conclude that the detrimental effect of the *Ec* R286Q mutation is due to the destabilization of the transducer rigidity affecting the allosteric transmission of ATP hydrolysis, rather than a direct effect due to alteration of DNA contact as previously suggested ^35^. Indeed the mutation of *Tth* K284 pointing in the cavity, and that could potentially mediate such interaction, has no effect on the catalytic activities.

### Analysis of the DNA-gate conformations

The deconvolution of different particles population in our dataset has led to distinct 3D reconstructions of the DNA-binding and cleavage domain. The 4.0 Å (State 1) and 4.6 Å (State 2) structures of the DNA-binding and cleavage domain bound to the double nicked G-segment reside in a configuration similar, but not identical, to the cleavage complex in presence of DNA and ciprofloxacin with a RMSD of 1.78 Å and 2.01 Å, respectively ^12^. The overall RMSD between State 1 and State 2 conformations is of 2.2 Å. In state 2, very subtle changes occur in structural elements lining the G-segment groove compared to State 1. The 2 conformations mostly differ in the GyrB subunit, more particularly the TOPRIM insertion that rotates around the GyrA N-terminal arm with a 5.0 Å upward displacement (Fig. 6a). This motion increases the distance between GyrA Tower and GyrB TOPRIM from 8 Å to 11 Å, inducing a slight shortening of the distance between the catalytic tyrosines from 26.4 Å to 25.6 Å and inducing the stretching of the G-segment by 2.5 Å in both directions (Fig. 6b and Supplementary Figure 10).

State 1 can be described as a cleavage complex in a closed conformation of the DNA gate, hereafter designated as ‘closed’. State 2 corresponds to an intermediate opening of the DNA-gate therefore referred as a ‘pre-opening’ conformation, when compared with the larger opening that can be observed in the conformation adopted by the hsTop2β ^36^ (Supplementary Figure 10d). State 1 and 2 correspond to a conformational oscillation before the opening of the DNA gate. This pre-opening transition is mainly undertaken by the insertion domain acting like an anvil which positions the TOPRIM domain in a configuration ready to perform the opening of the DNA-gate.

### High resolution features in the 4.0 Å structure of the Gyrase DNA-binding domain

Despite the high flexibility of the complex, we were able to solve a structure of the Gyrase DNA-binding domain in the closed complex with DNA and Gepotidacin at 4.0 Å resolution using a focused refinement strategy (Supplemental Figure 3). Side chains of secondary structure elements and the DNA double helix are very well defined in this highly ordered region (Fig. 7a). The local resolution near the NBTI binding site was estimated at 3 Å which allowed the identification of the Gepotidacin density (Fig. 7a-b and Supplementary Figure 4).

The molecule intercalates in the DNA duplex between positions +2 and +3 forming hydrophobic interactions and pi-stacks with the protein and the DNA, at the same location as previously observed for the NBTIs and crystallized with *S. aureus* DNA Gyrase ^21,24,37,38^ (Supplemental Figure 11a). The main difference in the binding pocket of *S. aureus* and *E. coli* gyrase lies in the subunit Methionine residue (M75) that is replaced by an Isoleucine residue in (I74) (Supplementary Figure S11a). This position is part of a GyrA hydrophobic pocket that accommodates the pyranopyridine ring of the Gepotidacin. Although the overall resolution of the pre-opening state is lower, a density for the Gepotidacin molecule could be identified at a lower rmsd level, suggesting that the Gepotidacin binding site is compatinble with conformational fluctuations at the DNA gate. Such flexibility of the DNA binding and cleavage domain in presence of Gepotidacin has also been observed in the *S. aureus* DNA crystal structures of DNA binding domains with an intact or doubly nicked DNA ^24^.

Further away along the DNA in the TOPRIM domain, a density could be observed in the 4 Å EM map at 6 Å rmsd at the cleavage reaction B-site that could correspond to the position of a magnesium ion (Supplementary Figure 12). The density of the conserved Aspartic and Glutamic residues defining the B-site are however much weaker but it has now been observed in high resolution cryo-EM maps that negatively charged groups yield weaker densities than positively charged groups ^39^. Taken together, it is likely that the cryo-EM structure could contain the requisite metal ions needed by DNA gyrase to perform DNA cleavage.

## Discussion

### Structural comparison of the *E. coli* and *T. thermophilus* DNA Gyrase DNA complex conformations

In both *T. thermophilus* and *E. coli* DNA gyrase complexes with DNA, 130 bp out of respectively 155 bp or 180 bp could be built, showing that the remaining 25 or 50 bp distributed on both sides of the G-segment are highly flexible ^12^. The spatial arrangement of the β-pinwheels induces an overall ∼150° bending of the G-segment in the *E. coli* Gyrase but only ∼75° in the *T. thermophilus* Gyrase (Fig. 4a). In addition, the β-pinwheels of both structures are swiveled by a 90° rotation angle in respect to each other (Fig. 4b). Interestingly, the 130 bp DNA duplex is chirally wrapped around the β-pinwheel orienting, in both structures, the T-segment in the DNA-gate groove formed by the TOPRIM and tower domains to form a 60° angle positive crossover when T-segment path is extrapolated. However, the angle of the T-segment entering the Gyrase core is different between the two species. In the *E. coli* structure, the combination of the high bending angle of the G-segment (∼150°) with the peculiar orientation of the β-pinwheels positions the T-segment to access the DNA-gate with an almost null or negative angle of ∼-10° (Fig. 4a). As a consequence, this configuration seems unfavorable to T-segment strand passage and would require a structural rearrangement of the β-pinwheel to proceed further. In contrast, the low bending angle of the G-segment (∼75°) and the position of the β-pinwheels in the *T. thermophilus* DNA Gyrase create a positive angle (10°) for the T-segment accessing the DNA-gate. This configuration seems more favorable to the T-segment strand passage without a structural rearrangement of the β-pinwheel.

Thus, both structures may be in two overall different states. The cryo-EM structures of the *T. thermophilus* and *E. coli* DNA gyrase complexes were determined in presence of ciprofloxacin and NBTI respectively that could have trapped two different conformational intermediates. The *E. coli* structure is reminiscent of a state immediately following G-segment binding and wrapping, preceding the conformation adopted by the *T. thermophilus* structure. Upon rotation of the β-pinwheels by 90°, the *T. thermophilus* structure would represent a state ready for T-segment passage. Interestingly, a recent single-molecule study described a Ω state, possibly similar to the *E. coli* complex and occurring before the α state in the catalytic cycle, more reminiscent of the *T. thermophilus* conformation ^13^. As previously suggested, such motion of the CTD in *E. coli* might be controlled and regulated by the acidic CTD tail in contact with the DNA-gate ^40^.

### Deconvolution of conformations and allosteric control of DNA Gyrase

During T-segment strand passage, the DNA-binding and cleavage domain needs to undergo several conformational changes. For decades, the opening of the DNA-gate was predicted to follow a “book-opening” motion, in which the two halves of the DNA-gate simply remain along the axis of G-segment ^2,3,41^. However, it has been shown recently that the DNA-gate opening mechanism is more subtle and is achieved by a “sliding and swiveling” motion of the two halves against each other, breaking the G-segment axis ^36^. The structure of the hsTop2β DNA-binding and cleavage domain provided by the aforementioned study is in an open conformation, revealing a funnel-shaped channel ready for the entry of T-segment. Besides, MD simulation of the T-segment crossing through the open DNA-gate suggested the presence of a conformation where the central cavity is widened and flattened, a state previously observed in a crystal structure of the *B. subtilis* DNA Gyrase ^42^. Consequently, the high degree of structural conservation of the bacterial and the human Top2β suggests that this mechanism of “sliding and swiveling” can be extended to the bacterial Topo 2A, and thus to DNA Gyrase. In this context, the pre-opening conformation of the cleavage complex represents one of the discreet steps that DNA Gyrase undergoes to open the DNA-gate (Supplementary Movie 2).

In *E. coli*, this step might be further controlled by the presence of the GyrB insertion domain. Deletion of the insertion was shown to greatly reduce the DNA binding, supercoiling and DNA-stimulated ATPase activities of *E. coli* DNA Gyrase ^17^. The subtle swiveling movement of the GyrB insertion domain around the N-terminal arm of GyrA leads to a controlled stretching of the G-segment by 5 Å and spatial separation of the TOPRIM and Tower domains, a necessary step for the formation of the groove that will accommodate the T-segment. In the light of our structural study, the insertion in the TOPRIM domain may presumably act as a steric counterweight amplifying the movements of the DNA-gate.

The DNA gate opening is coupled to ATP hydrolysis through the transducer region whose cavity is thought to accommodate a DNA T-segment, as evidenced by recent structural data on a bacterial Topo IV ATPase domain with an oligonucleotide ^43^. Basic residues lining the transducer cavity and in particular residue R286 in the *E. coli* ATPase domain are thought to contribute to DNA capture and enzyme allostery ^35^. The GHKL region of the ATPase domain shows a strong sequence identity across species due to the universal conservation of the catalytic motifs, with more sequence variability across the transducer sequence (Supplementary Figure 8). Sequence alignments show that, at this position of the transducer helix, gyrase B possesses an arginine like *E. coli* or is replaced by a lysine in other organisms, such as the *Thermus* homolog. Although the positive charge of the side chains may be compatible with a common role in DNA binding and transport, point mutation of the arginine in the *α*-helix of the transducer has a different impact than mutation of the lysine at this same position, suggesting a specific catalytic regulation in *E. coli* through this residue. However, the analysis of the structures shows that *E. coli* R286 is engaged in a salt bridge stabilizing the transducer structure, a feature that is replaced by a hydrophobic core in the *Thermus* enzyme, rather indicative of a structural element maintaining transducer rigidity. It is not clear at this stage if DNA capture and transport is directly linked to the basic residues in the transducer and at which catalytic step molecular interaction with the T-segment intervenes. The full architecture of the complex in presence of a T-segment still needs to be determined to better understand the molecular determinants of DNA capture. Despite the structural specificities of the *E. coli* enzyme, this demonstrates that the rigidity of the transducer region is essential for the enzyme activities and efficient allosteric transmission.

### Position of the GyrA box and β-pinwheel

The complete DNA binding around the β-pinwheel surface generates a twisting of ∼200° consistent with the reported wrapping around DNA gyrase pinwheel blades ^44^. The highly conserved GyrA box motif, QRRGGKG ^45^, located on the first blade of the β-pinwheels is essential for DNA wrapping (Fig. 3a-b). Complete deletion or alanine substitution of this motif completely abolishes the negative supercoiling activity ^46^. By elucidating the spatial localization of the GyrA box motif when DNA Gyrase is bound to DNA, we show here why the GyrA box motif is crucial in wrapping DNA. The position of the GyrA box suggests that it acts as a clamp maintaining the DNA wrapped around the β-pinwheel with strong interaction that can compensate for the energetically unfavorable configuration of the twisted DNA (Fig. 3d). This explains the role of the GyrA box behaving as a ‘at the end of the wrap’ mechanism, consistent with previous studies ^47,48^.

In addition, the analysis of the complex overall conformation shows that the β-pinwheels are oriented in a way that the CTD tail could face the GyrB TOPRIM domain. The CTD sequence of *E. coli* DNA gyrase is composed of an unfolded and poorly conserved acidic tail starting at the end of the β-pinwheel structure (Supplementary Figure 13a). Several biochemical studies have shown that the CTD tail stimulates DNA wrapping coupled with ATP hydrolysis ^40,49^. Modeling of the 34 residues of the *E. coli* CTD tail shows that it could easily reach and contact the TOPRIM insertion domain (Supplementary Figure 13b). Conformational changes of the DNA-gate from closed to pre-opening state may conduct a signal to the β-pinwheel through the tethered acidic tail promoting a β-pinwheel vertical movement. This interaction might also drive the relative orientation of the β-pinwheels wrapped with DNA, positioning the T-segment in the proper direction for transport to the DNA-gate.

### High resolution features and drug binding site

The high resolution features of the DNA binding domain structure now makes it possible to detect the density of small molecules and possibly catalytic ions in the cryo-EM map. We were able to observe the Gepotidacin molecule inserted in the DNA a on the two-fold axis at the GyrA dimer interface, as previously observed in crystal structures of the *S. aureus* DNA Gyrase in complex with the NBTI ^21,24,37,38^. Gepotidacin is currently in clinical trials for the treatment of *N. gonorrhoeae* infections ^22,23^, a species that is phylogenetically closer to *E. coli* and also harbors an Isoleucine at this position in the Gyrase sequence (Supplementary Figure 11b-c). The comparison of the cryo-EM map with the electron density map of crystal structures ^24^ shows that the cryo-EM map information is of comparable quality in this region for the identification of small compounds (Supplementary Figure 14).

Based on this study, we foresee that it is now possible to consider cryo-EM as a viable tool to perform structure-guided drug design to target flexible complexes and identify new conformations of DNA Gyrase that could so far not be obtained by conventional structural methods.

## Methods

### GyrB and GyrA expression and purification

The sequence coding for the full-length *E. coli* GyrA (2-875) was inserted into a modified pET28b containing an N-terminal 10-His tag and a C-terminal Twin-strep tag. Overexpression was performed in *E. coli* BL21 (DE3) pRARE2 in LB medium containing 50 µL/L kanamycin and 35 µL/L chloramphenicol. Cells were induced with 0.35 mM IPTG after reaching an OD_600_0.85 and protein was expressed at 37°C for 4 h. Cells were harvested and resuspended in lysis buffer (20 mM Hepes, 500 mM NaCl, 20 mM imidazole, 10% v/v glycerol, pH 8.0) and lysed with 3 cycles of high-pressure disruption using EmulsiFlex-C3 at 1500 bars. The GyrA protein was purified by nickel-affinity chromatography on a manually packed XK 26/20 column (Pharmacia) with Chelating Sepharose 6 Fast Flow resin (GE Healthcare) bound to Ni^2+^ ions. Elution was performed with the lysis buffer containing 250 mM imidazol pH 8.0 and eluted proteins were directly attached on a 10 ml Streptavidin Sepharose (GE Healthcare). Proteins were extensively washed with Strep buffer (20 mM Hepes, 60 mM NaCl, 1 mM EDTA, 1 mM DTT, 10% v/v glycerol, pH 8.0) and eluted with Strep buffer containing 3 mM Desthiobiotin (Sigma-Aldrich). Both 10-His tag and Twin-strep tag were subsequently cleaved by PreScission protease (P3C) and Tobacco Etch Virus (TEV) cleavage (mass ratio 1:1:50 P3C-TEV-GyrA) overnight at 4°C. GyrA was then further purified by an anion exchange chromatography step using a HiTrap Q HP column (GE Healthcare). The protein was eluted with a linear gradient of 20 column volumes with buffer B (20 mM Hepes, 1 M NaCl, 1 mM EDTA, 1 mM DTT, 10% v/v glycerol, pH 8.0). Fractions containing GyrA were pooled and loaded on a Superdex S200 16/60 size exclusion chromatography column (GE Healthcare) using 20 mM Hepes, 50 mM NaGlu, 50 mM KAc, 1 mM EDTA, 0.5 mM DTT, 10% v/v glycerol, pH 8.0. 37 mg of GyrA were obtained from 3 L of cultures. The GyrB (2-804) coding sequence was inserted into the same modified pET28b. Overexpression was performed in *E. coli* BL21 (DE3) pRARE2 in LB medium containing 50 µL/L kanamycin and 35 µL/L chloramphenicol. Cells were induced with 0.35 mM IPTG after reaching an OD_600_ 0.85 and protein was expressed at 18°C for 18 h. The GyrB purification procedure is as described above for GyrA. 10 mg of GyrB were obtained from 3 L of cultures.

### Full DNA Gyrase reconstitution

*E. coli* GyrA and GyrB were mixed at equimolar ratio to allow full DNA Gyrase reconstitution. The complex was further purified on a Superdex S200 16/60 size exclusion chromatography column (GE Healthcare) using cryo-EM buffer (20 mM Hepes, 30 mM NaGlu, 30 mM KAc, 2 mM MgAc, 0.5 mM TCEP, pH 8.0) (Fig. 1 and Supplementary Figure 1).

### Nucleic acid preparation

A double nicked 180bp DNA duplex was reconstituted using 2 phosphorylated asymmetric synthetic oligonucleotides ^12^ obtained from Sigma-Aldrich (Supplemental Table 1). Briefly, the nucleic acids were dissolved in DNAse-free water at 1 mM concentration. To assemble the double stranded DNA, each oligo was mixed at 1:1 molar ratio, annealed by incubating at 95°C for 2 min and then decreasing the temperature by 1°C every 1 min until reaching 20°C. The annealed doubly nicked DNA duplex was then buffer-exchanged in Hepes 20 mM pH 8.0 with a BioSpin 6 column (BioRad).

### Complex formation for cryo-EM

The purified DNA Gyrase was mixed with the 180 bp dsDNA at 1:1 molar ratio with a final protein and DNA concentration of 32 µM. The mixture was incubated for 10 min at 37°C. Gepotidacin (GSK 2140944) resuspended at 10 mM in 100% DMSO was added to reach a final concentration of 170 µM (1.7% DMSO). The mixture was incubated for 10 min at 37°C. AMP-PNP (Sigma) was added to the complex at a final concentration of 340 µM. The fully reconstituted complex was further incubated 30 min at 30°C. Finally, 8 mM CHAPSO (Sigma-Aldrich) was added to the complex. The sample was centrifuged 2 h at 16,000 g to remove potential aggregates.

### Cryo-EM grid preparation

Quantifoil R-1.2/1.3 300 mesh copper grids were glow-charged for 20 s prior to the application of 4 µl of the complex. After 30 s of incubation, the grids were plunge-frozen in liquid ethane using a Vitrobot mark IV (FEI) with 95% chamber humidity at 10°C.

### Electron microscopy

Cryo-EM imaging was performed on a Titan Krios microscope operated at 300 kV (FEI) equipped with a K2 Summit direct electron camera (Gatan), a GIF Quantum energy filter (Gatan) operated in zero-energy-loss mode with a slit width of 20 e^-^V, a Volta Phase-Plate (FEI) and a CS corrector (FEI). Images were recorded in EFTEM nanoprobe mode with Serial EM ^50^ in super-resolution counting mode at nominal magnification of 130,000x with a super resolution pixel size of 0.44 Å and a constant defocus target of −500 nm (Supplementary Figure 2c). The VPP was advanced to a new position every 100 min (Supplementary Figure 2b). Four datasets were collected with a dose rate of 6 to 8 e^-^/pixel/s (0.88 Å pixel size at the specimen) on the detector. Images were recorded with a total dose of 50 e^-^/Å^2^, exposure time between 7 to 10 s and 0.2 to 0.25 s subframes (35 to 50 total frames). A total of 11,833 movies were recorded after data collection of the 4 datasets.

### Data processing

Data processing of each data set was done separately following the same procedure until the 3D refinements when particles were merged. The super-resolution dose-fractionated subframes were gain-corrected with IMOD ^51^ and binned twice by Fourier-cropping, drift-corrected and dose-weighted using MotionCor2 ^52^ yielding summed images with 0.88 Å pixel size. The contrast transfer function of the corrected micrographs was estimated using GCTF v1.06 ^53^. Thon rings were manually inspected for astigmatism and micrographs with measured resolutions worse than 4 Å were discarded yielding 8,701 remaining micrographs. Particles were automatically picked by reference-free Gaussian blob picking protocol in RELION2 ^28,29^. Taking together the 4 data sets, a total of 1,572,962 particles were selected. Four times binned particles from each data set were separately subjected to 2 rounds of 2D classification in RELION2 to remove junk particles and contaminations resulting in a total of 479,667 particles for further processing. Particles from the 4 data sets were merged into one unique data set followed by a re-extraction with centering in 3×3 binned format. A last round of 2D classification was then performed yielding a final particle stack of 338,616 particles. The subsequent particle stack was subjected to one round of 3D *ab-initio* classification in cryoSPARC ^30^. After discarding the poor-quality models, the remaining ab-initio model resulted in a final data set of 191,456 particles, with a class probability threshold of 0.9 (Supplementary Figure 2a). The *ab-initio* model was low-pass filtered to 30 Å and was used as a reference for homogeneous refinement in cryoSPARC resulting in a 5.4 Å map. The refined particle coordinates were then used for local CTF estimation using GCTF v1.06 followed by re-extraction of 2×2 binned particles with centering. This new particle stack was subjected to a 3D auto-refinement in RELION2 using the *ab-initio* model low-pass filtered at 30 Å yielding a map with a global resolution of 5.0 Å (Supplementary Figure 3). Local resolution showed a range of resolution from 4 Å in the DNA-binding core to 9 Å in the ATPase head and the GyrA β-pinwheel wrapping the DNA, indicative of high flexibility of these two modules. To obtain a final reconstruction of the full complex with well-defined densities for each domain, we performed 3D classification without alignment yielding several different classes. Particles from 3 classes were merged (94,633 particles). The subsequent 1×1-binned particles stack was refined in RELION2 without mask until late refinement iterations where a soft mask was applied to improve resolution. Post-processing of the map yielded a 6.6 Å resolution reconstruction of the overall complex (Supplementary Figure 3).

A combination of different local approaches was used to identify different conformations and to improve local resolution of each domain. Since the ATPase head domain was too small for a 3D focused refinement, we first performed a focused 3D classification of the ATPase head with a soft mask and no alignment in RELION2. One class of 58,329 particles with a well-defined density of the ATPase head was selected, followed by re-extraction of 1×1-binned particles yielding particles with pixel size of 0.88 Å/pixel. To facilitate an accurate alignment of the particles, this class was further refined in RELION2 using a soft mask around the ATPase head and the DNA-binding core yielding a 5.9 Å global resolution map (ATPase-Core) (Supplementary Figure 3).

Secondly, we performed a focused 3D classification of the GyrA β-pinwheel with a soft mask and no alignment in RELION2. One class of 45,040 particles with a well-defined density of the GyrA β-pinwheel was selected, followed by re-extraction of 1×1-binned particles yielding particles with pixel size of 0.88 Å/pixel. This class was further refined in RELION2 using a soft mask around the β-pinwheel and the DNA-binding core yielding a 6.3 Å resolution map (CTD-Core) (Supplementary Figure 3).

Finally, a focused 3D auto-refinement was performed in RELION2 using a soft mask around the DNA-binding core. Post-processing of the map produced a 4.3 Å resolution reconstruction of the DNA-binding core. Then, a focused 3D classification of the DNA-binding was performed. Two of the classes showed better angular accuracies and distinct “Closed” (60,548 particles) and “Pre-opening” conformations (53,655 particles) of the DNA-binding core. These 2 classes were further refined in RELION2 by focused 3D refinement with a C2 symmetry using 1×1 binned particles and gave reconstructions with global resolution of 4.0 Å and 4.6 Å after post-processing, respectively (Supplementary Figure 3). The “Closed” DNA-binding core was further refined by focused 3D refinement using a soft mask which excludes the disordered TOPRIM insertion yielding a 4.0 Å map of high quality (Supplementary Figure 3).

All reported resolutions are based on the gold standard FSC-0.143 criterion ^54^ and FSC-curves were corrected for the convolution effects of a soft mask using high-resolution noise-substitution ^55^ in RELION2 as well as in cryoSPARC (Supplementary Figure 4a). All reconstructions were sharpened by applying a negative B-factor that was estimated using automated procedures ^56^. Local resolution of maps was calculated using Blocres ^57^ (Supplementary Figure 4c). The cryo-EM maps of the overall complex, ATPase-Core, CTD-Core and the DNA-binding and cleavage domain in closed (with and without TOPRIM insertion) and in pre-opening state have been deposited in the EM Data Bank under accession numbers EMD-4913, EMD-4914, EMD-4915, EMD-4910, EMD-4909, EMD-4912, respectively.

### Model building and refinement of the DNA-binding core

The EM reconstruction of the DNA-binding core lacking the TOPRIM insertion solved at 4.0 Å was of the best quality. Almost all side chains could be seen and Gepotidacin density was clearly visible. This map was used to refine a crystal structure of the DNA binding core of the *E. coli* DNA Gyrase ^17^. A dimeric atomic model of the DNA-binding core was generated using PDB 3NUH in PyMol (Schrodinger L.L.C.). The subsequent atomic model was stripped of amino acids belonging to the TOPRIM insertion, all ions and water molecules, with all occupancies set to 1 and B-factors set to 50. First, the atomic model was rigid-body fitted in the filtered and sharpened map with Chimera ^58^. A first round of real-space refinement in PHENIX ^59^ was performed using local real-space fitting and global gradient-driven minimization refinement. Then, 20 nucleic acids from the structure *of S*.*aureus* DNA gyrase ^38^ (PDB 5IWM) were copied and fitted into our atomic model. DNA sequence was modified according to the DNA used in our structure. Missing protein residues and nucleic acids were manually built in COOT ^60^. Atomic model and constraint dictionary of Gepotidacin (GSK-2140944) were generated with the Grade server (http://grade.globalphasing.org). Gepotidacin was then manually fitted in the empty electron density in COOT and then duplicated. Both duplicates were set to 0.5 occupancy. Several rounds of real-space refinement in PHENIX using restraints for secondary structure, rotamers, Ramachandran, and non-crystallographic symmetry were performed, always followed by manual inspection in COOT, until a converging model was obtained. Finally, B-factors were refined by a final round of real-space refinement in PHENIX using the same settings as before. After the refinement has converged, atomic coordinates of the TOPRIM insertion were added to the atomic model. It was further refined in the EM reconstructions containing the insertion density in closed and pre-open states using the same procedure. A half-map cross-validation was performed to define 1.5 as the best refinement weight in PHENIX allowing atom clash reduction and prevention of model overfitting (Supplementary Figure 4e). All refinement steps were done using the resolution limit of the reconstructions according to the gold standard FSC-0.143 criterion ^54^. Refinement parameters, model statistics and validation scores are summarized in Supplementary Table 2. The atomic models of the DNA-binding and cleavage domain in closed (with and without insertion) and pre-opening conformations have been deposited in the Protein Data Bank under accession numbers 6RKU, 6RKS, 6RKV, respectively.

### Model building and refinement of the overall complex

We used a combination of different EM reconstructions to build the DNA Gyrase overall complex atomic model. The “ATPase-Core” structure solved at 5.9 Å was used to rigid body fit in Chimera the ATPase domain crystal structure in complex with ADPNP ^31^ (PDB 1EI1) and the DNA-binding core refined in closed conformation. The quality of the electron density allowed to build in COOT the missing residues of the linker between the ATPase domain and the DNA-binding and cleavage domain. The “CTD-Core” structure solved at 6.3 Å was used to accurately rigid body fit the β-pinwheel crystal structure ^10^ in Chimera. The 10 missing amino acids (564-574) following the GyrA-box were added using the modelling server Phyre2 ^61^. The quality of the electronic density allowed to build in COOT the missing nucleic acids around the β-pinwheel as well as the 10 amino acids linking the last GyrA residues to the β-pinwheel. Based on a secondary structure prediction, a part of the linker was built as an alpha helix (Fig. 3). The subsequent atomic model containing the ATPase domain, the DNA-binding core and one β-pinwheel was rigid body fitted into the overall complex structure solved at 6.6 Å using Chimera. Then, a copy of the first β-pinwheel was fitted into the density of the second β-pinwheel in COOT. The missing nucleic acids around the second β-pinwheel as well as the missing 10 residues of the linker were manually built in COOT. The resulting atomic model was stripped of all ions and water molecules, with all occupancies set to 1 and B-factors set to 50. Finally, real-space refinement of the atomic model against the overall complex structure was performed in PHENIX using rigid-body and global gradient-driven minimization refinement. Resolution limit for refinement was set according to the gold standard FSC-0.143 criterion ^54^. Half-map cross-validations were also performed (Supplementary Figure 4e). Refinement parameters, model statistics and validation scores are summarized in Supplementary Table 2. The atomic model of the overall structure has been deposited in the Protein Data Bank under accession numbers 6RKW. All the Figures were created with Chimera ^58^, ChimeraX ^62^ and PyMol (Schrodinger L.L.C.).

### *E. coli* GyrB R286K, R286Q expression and purification

The modified pET28b used for wild-type *E. coli* GyrB overexpression was mutated by site directed mutagenesis using the QuikChange XL Site-Directed Mutagenesis kit (Agilent) in order to generate two plasmids harboring R286K or R286Q mutations. Overexpression and purification procedures for the two mutants are identical to the wild-type GyrB described above in the Methods section.

### *T. thermophilus* GyrB K284R, K284Q expression and purification

The modified pET28b used for wild-type *T. thermophilus* GyrB overexpression ^12^ was mutated by site directed mutagenesis using the QuikChange XL Site-Directed Mutagenesis kit (Agilent) in order to generate two plasmids harboring K284R or K284Q mutations. Overexpression and purification procedures for the two mutants are identical to the *E. coli* wild-type GyrB described above in the Methods section.

### DNA supercoiling assay

An increasing concentration of DNA Gyrase (GyrA_2_B_2_) was incubated at 37°C with 6 nmoles of relaxed pUC19 plasmid in a reaction mixture containing 20 mM Tris-acetate pH 7.9, 100 mM potassium acetate, 10 mM magnesium acetate, 1 mM DTT, 1 mM ATP, 100 µg/ml BSA. After 30 minutes, reactions were stopped by addition of SDS 1%. Agarose gel electrophoresis was used to monitor the conversion of relaxed pUC19 to the supercoiled form. Samples were run on a 0.8% agarose, 1X Tris Borate EDTA buffer (TBE) gel, at 6 V/cm for 180 min at room temperature. Agarose gels were stained with 0.5 mg/ml ethidium bromide in 1X TBE for 15 min, followed by 5 min destaining in water. DNA topoisomers were revealed using a Typhoon (GE Healthcare).

### ATPase activity assay

ATPase activity assays were performed as described in ^63^. ATP hydrolysis is measured by following the oxidation of NADH mediated by pyruvate kinase (PK) and lactate dehydrogenase (LDH). The absorbance was monitored at 340 nm over 600 seconds at 37°C with a Shimadzu 1700 spectrophotometer. Reactions were recorded in triplicates with 100 nM of GyrA_2_B_2_ and 16 nM linear DNA (pCR-blunt) in 500 µl of a buffer containing 50 mM Tris HCl pH 7.5, 150 mM potassium acetate, 8 mM magnesium acetate, 7 mM BME, 100 µg/mg of BSA, 4U/5U of PK/LDH mixture, 2 mM PEP, and 0.2 mM NADH.

### Multiple alignment, evolutionary conservation of residues and phylogenetic tree

GyrB ATPase/Transducer protein sequences from 30 species (Uniprot codes: GYRB_STRCO, *Streptomyces coelicolor*; A0A3A2T2C3_LISMN, *Listeria monocytogenes*; W8F4Q2_AGRT4, *Agrobacterium tumefasciens*; Q83FD5_COXBU, Coxiella burnetii; GYRB_ECOLI, *Escherichia coli*; GYRB_NEIGO, *Neisseria gonorrhoeae*; GYRB_SALTY, *Salmonella typhimurium*; Q7V9N3_PROMA, *Prochlorococcus marinus*; GYRB_STRR6, *Streptococcus pneumoniae*; GYRB_MYCPN, *Mycoplasma pneumoniae*; GYRB_SHIFL, Shigella flexneri; GYRB_PSEAE, *Pseudomonas aeruginosa*, Q18C89_PEPD6, *Clostridium difficile*; GYRB_RICFE, *Rickettsia felis*; Q8CZG1_YERPE, *Yersinia pestis*; GYRB_BORBU; *Borrelia burgdorferi*; GYRB_ENTFA, *Enterococcus faecalis*; Q8YEQ5_BRUME, *Brucella melitensis*; GYRB_STAAU, *Staphylococcus aureus*; Q7VQU8_BLOFL, *Blochmannia floridanus*; GYRB_HAEIN, *Haemophilus influenzae*; GYRB_MYCTU, *Mycobacterium tuberculosis*, GYRB_THEMA, *Thermotoga maritima*; GYRB_THET8, *Thermus thermophilus*; GYRB_BACSU, *Bacillus subtilis*; GYRB_HELPY, *Helicobacter pylori*; GYRB_CHLTR, *Chlamydia trachomatis*; GYRB_VIBCH, *Vibrio cholerae*; GYRB_THEAQ, *Thermus aquaticus*; GYRB_AQUAE, *Aquifex aeolicus*) were aligned using the Clustal Omega server (EMBL-EBI). The subsequent alignment was used to plot the amino acids evolutionary conservation on the ATPase/transducer structure (PDB ID 1EI1) using the ConSurf server (http://consurf.tau.ac.il). The phylogenetic tree was generated by neighbour-joining method using the multiple alignment in Clustal Omega.

## Data availability

Model coordinates and density maps are available in the Protein Data Bank (PDB ID 6RKS, 6RKU, 6RKV, 6RKW) and the EM Data Bank (EMD-4909, EMD-4910, EMD-4912, EMD-

4913, EMD-4914, EMD-4915).

## Acknowledgements

We thank the teams of B. Klaholz and P. Schultz, as well as A. Weixlbaumer and X. Guo for useful technical suggestions and help with computational resources. This work was supported by the Fondation ARC and the grant ANR-10-LABX-0030-INRT (managed by the Agence Nationale de la Recherche under the frame programme Investissements d’Avenir ANR-10-IDEX-0002-02). The authors acknowledge the support and the use of resources of the French Infrastructure for Integrated Structural Biology FRISBI ANR-10-INBS-05 and of Instruct-ERIC. Computational resources were provided by the Méso-centre de Calcul (University of Strasbourg).

## Contributions

A.V.B. and V.L. conceived the study and designed the experiments. A.V.B and J.O. performed the experiments. A.V.B. and V.L. analyzed and interpreted the data. A.V.B. and V.L. wrote the manuscript.

## Competing interests

The authors declare no competing interests.

## References

1. Wang, J.C. Cellular roles of DNA topoisomerases: a molecular perspective. Nat Rev Mol Cell Biol 3, 430–40 (2002).

2. Vos, S.M., Tretter, E.M., Schmidt, B.H. & Berger, J.M. All tangled up: how cells direct, manage and exploit topoisomerase function. Nat Rev Mol Cell Biol 12, 827–41 (2011).

3. Schoeffler, A.J. & Berger, J.M. DNA topoisomerases: harnessing and constraining energy to govern chromosome topology. Q Rev Biophys 41, 41–101 (2008).

4. Bush, N.G., Evans-Roberts, K. & Maxwell, A. DNA Topoisomerases. EcoSal Plus 6(2015).

5. Gellert, M., Mizuuchi, K., O’Dea, M.H. & Nash, H.A. DNA gyrase: an enzyme that introduces superhelical turns into DNA. Proc Natl Acad Sci U S A 73, 3872–6 (1976).

6. Roca, J. & Wang, J.C. DNA transport by a type II DNA topoisomerase: evidence in favor of a two-gate mechanism. Cell 77, 609–16 (1994).

7. Roca, J., Berger, J.M., Harrison, S.C. & Wang, J.C. DNA transport by a type II topoisomerase: direct evidence for a two-gate mechanism. Proc Natl Acad Sci U S A 93, 4057–62 (1996).

8. Wigley, D.B., Davies, G.J., Dodson, E.J., Maxwell, A. & Dodson, G. Crystal structure of an N-terminal fragment of the DNA gyrase B protein. Nature 351, 624–9 (1991).

9. Morais Cabral, J.H. et al. Crystal structure of the breakage-reunion domain of DNA gyrase. Nature 388, 903–6 (1997).

10. Ruthenburg, A.J., Graybosch, D.M., Huetsch, J.C. & Verdine, G.L. A superhelical spiral in the Escherichia coli DNA gyrase A C-terminal domain imparts unidirectional supercoiling bias. J Biol Chem 280, 26177–84 (2005).

11. Soczek, K.M., Grant, T., Rosenthal, P.B. & Mondragon, A. CryoEM structures of open dimers of gyrase A in complex with DNA illuminate mechanism of strand passage. Elife 7(2018).

12. Papillon, J. et al. Structural insight into negative DNA supercoiling by DNA gyrase, a bacterial type 2A DNA topoisomerase. Nucleic Acids Res 41, 7815–27 (2013).

13. Basu, A. et al. Dynamic coupling between conformations and nucleotide states in DNA gyrase. Nat Chem Biol 14, 565–574 (2018).

14. Basu, A., Parente, A.C. & Bryant, Z. Structural Dynamics and Mechanochemical Coupling in DNA Gyrase. J Mol Biol 428, 1833–45 (2016).

15. Gubaev, A. & Klostermeier, D. The mechanism of negative DNA supercoiling: a cascade of DNA-induced conformational changes prepares gyrase for strand passage. DNA Repair (Amst) 16, 23–34 (2014).

16. Stelljes, J.T., Weidlich, D., Gubaev, A. & Klostermeier, D. Gyrase containing a single C-terminal domain catalyzes negative supercoiling of DNA by decreasing the linking number in steps of two. Nucleic Acids Res 46, 6773–6784 (2018).

17. Schoeffler, A.J., May, A.P. & Berger, J.M. A domain insertion in Escherichia coli GyrB adopts a novel fold that plays a critical role in gyrase function. Nucleic Acids Res 38, 7830–44 (2010).

18. Collin, F., Karkare, S. & Maxwell, A. Exploiting bacterial DNA gyrase as a drug target: current state and perspectives. Appl Microbiol Biotechnol 92, 479–97 (2011).

19. Heide, L. New aminocoumarin antibiotics as gyrase inhibitors. Int J Med Microbiol 304, 31–6 (2014).

20. Delgado, J.L., Hsieh, C.M., Chan, N.L. & Hiasa, H. Topoisomerases as anticancer targets. Biochem J 475, 373–398 (2018).

21. Bax, B.D. et al. Type IIA topoisomerase inhibition by a new class of antibacterial agents. Nature 466, 935–40 (2010).

22. Biedenbach, D.J. et al. In Vitro Activity of Gepotidacin, a Novel Triazaacenaphthylene Bacterial Topoisomerase Inhibitor, against a Broad Spectrum of Bacterial Pathogens. Antimicrob Agents Chemother 60, 1918–23 (2016).

23. Scangarella-Oman, N.E. et al. Microbiological Analysis From a Phase 2 Randomized Study in Adults Evaluating Single Oral Doses of Gepotidacin in the Treatment of Uncomplicated Urogenital Gonorrhea Caused by Neisseria gonorrhoeae. Antimicrob Agents Chemother (2018).

24. Gibson, E.G., Bax, B., Chan, P.F. & Osheroff, N. Mechanistic and Structural Basis for the Actions of the Antibacterial Gepotidacin against Staphylococcus aureus Gyrase. ACS Infect Dis 5, 570–581 (2019).

25. Pommier, Y. Drugging topoisomerases: lessons and challenges. ACS Chem Biol 8, 82–95 (2013).

26. Danev, R. & Baumeister, W. Cryo-EM single particle analysis with the Volta phase plate. Elife 5(2016).

27. Khoshouei, M., Radjainia, M., Baumeister, W. & Danev, R. Cryo-EM structure of haemoglobin at 3.2 A determined with the Volta phase plate. Nat Commun 8, 16099 (2017).

28. Kimanius, D., Forsberg, B.O., Scheres, S.H. & Lindahl, E. Accelerated cryo-EM structure determination with parallelisation using GPUs in RELION-2. Elife 5(2016).

29. Scheres, S.H. RELION: implementation of a Bayesian approach to cryo-EM structure determination. J Struct Biol 180, 519–30 (2012).

30. Punjani, A., Rubinstein, J.L., Fleet, D.J. & Brubaker, M.A. cryoSPARC: algorithms for rapid unsupervised cryo-EM structure determination. Nat Methods 14, 290–296 (2017).

31. Brino, L. et al. Dimerization of Escherichia coli DNA-gyrase B provides a structural mechanism for activating the ATPase catalytic center. J Biol Chem 275, 9468–75 (2000).

32. Dong, K.C. & Berger, J.M. Structural basis for gate-DNA recognition and bending by type IIA topoisomerases. Nature 450, 1201–5 (2007).

33. Schmidt, B.H., Osheroff, N. & Berger, J.M. Structure of a topoisomerase II-DNA-nucleotide complex reveals a new control mechanism for ATPase activity. Nat Struct Mol Biol 19, 1147–54 (2012).

34. Stanger, F.V., Dehio, C. & Schirmer, T. Structure of the N-terminal Gyrase B fragment in complex with ADPPi reveals rigid-body motion induced by ATP hydrolysis. PLoS One 9, e107289 (2014).

35. Tingey, A.P. & Maxwell, A. Probing the role of the ATP-operated clamp in the strand-passage reaction of DNA gyrase. Nucleic Acids Res 24, 4868–73 (1996).

36. Chen, S.F. et al. Structural insights into the gating of DNA passage by the topoisomerase II DNA-gate. Nat Commun 9, 3085 (2018).

37. Chan, P.F. et al. Thiophene antibacterials that allosterically stabilize DNA-cleavage complexes with DNA gyrase. Proc Natl Acad Sci U S A 114, E4492–E4500 (2017).

38. Miles, T.J. et al. Novel tricyclics (e.g., GSK945237) as potent inhibitors of bacterial type IIA topoisomerases. Bioorg Med Chem Lett 26, 2464–2469 (2016).

39. Wang, J. & Moore, P.B. On the interpretation of electron microscopic maps of biological macromolecules. Protein Sci 26, 122–129 (2017).

40. Tretter, E.M. & Berger, J.M. Mechanisms for defining supercoiling set point of DNA gyrase orthologs: I. A nonconserved acidic C-terminal tail modulates Escherichia coli gyrase activity. J Biol Chem 287, 18636–44 (2012).

41. Wendorff, T.J., Schmidt, B.H., Heslop, P., Austin, C.A. & Berger, J.M. The structure of DNA-bound human topoisomerase II alpha: conformational mechanisms for coordinating inter-subunit interactions with DNA cleavage. J Mol Biol 424, 109–24 (2012).

42. Rudolph, M.G. & Klostermeier, D. Mapping the spectrum of conformational states of the DNA- and C-gates in Bacillus subtilis gyrase. J Mol Biol 425, 2632–40 (2013).

43. Laponogov, I. et al. Trapping of the transport-segment DNA by the ATPase domains of a type II topoisomerase. Nat Commun 9, 2579 (2018).

44. Corbett, K.D., Shultzaberger, R.K. & Berger, J.M. The C-terminal domain of DNA gyrase A adopts a DNA-bending beta-pinwheel fold. Proc Natl Acad Sci U S A 101, 7293–8 (2004).

45. Ward, D. & Newton, A. Requirement of topoisomerase IV parC and parE genes for cell cycle progression and developmental regulation in Caulobacter crescentus. Mol Microbiol 26, 897–910 (1997).

46. Kramlinger, V.M. & Hiasa, H. The “GyrA-box” is required for the ability of DNA gyrase to wrap DNA and catalyze the supercoiling reaction. J Biol Chem 281, 3738–42 (2006).

47. Costenaro, L., Grossmann, J.G., Ebel, C. & Maxwell, A. Small-angle X-ray scattering reveals the solution structure of the full-length DNA gyrase a subunit. Structure 13, 287–96 (2005).

48. Lanz, M.A. & Klostermeier, D. The GyrA-box determines the geometry of DNA bound to gyrase and couples DNA binding to the nucleotide cycle. Nucleic Acids Res 40, 10893–903 (2012).

49. Lanz, M.A., Farhat, M. & Klostermeier, D. The acidic C-terminal tail of the GyrA subunit moderates the DNA supercoiling activity of Bacillus subtilis gyrase. J Biol Chem 289, 12275–85 (2014).

50. Mastronarde, D.N. Automated electron microscope tomography using robust prediction of specimen movements. J Struct Biol 152, 36–51 (2005).

51. Mastronarde, D.N. & Held, S.R. Automated tilt series alignment and tomographic reconstruction in IMOD. J Struct Biol 197, 102–113 (2017).

52. Zheng, S.Q. et al. MotionCor2: anisotropic correction of beam-induced motion for improved cryo-electron microscopy. Nat Methods 14, 331–332 (2017).

53. Zhang, K. Gctf: Real-time CTF determination and correction. J Struct Biol 193, 1–12 (2016).

54. Scheres, S.H. & Chen, S. Prevention of overfitting in cryo-EM structure determination. Nat Methods 9, 853–4 (2012).

55. Chen, S. et al. High-resolution noise substitution to measure overfitting and validate resolution in 3D structure determination by single particle electron cryomicroscopy. Ultramicroscopy 135, 24–35 (2013).

56. Rosenthal, P.B. & Henderson, R. Optimal determination of particle orientation, absolute hand, and contrast loss in single-particle electron cryomicroscopy. J Mol Biol 333, 721–45 (2003).

57. Heymann, J.B. & Belnap, D.M. Bsoft: image processing and molecular modeling for electron microscopy. J Struct Biol 157, 3–18 (2007).

58. Pettersen, E.F. et al. UCSF Chimera--a visualization system for exploratory research and analysis. J Comput Chem 25, 1605–12 (2004).

59. Adams, P.D. et al. PHENIX: a comprehensive Python-based system for macromolecular structure solution. Acta Crystallogr D Biol Crystallogr 66, 213–21 (2010).

60. Emsley, P., Lohkamp, B., Scott, W.G. & Cowtan, K. Features and development of Coot. Acta Crystallogr D Biol Crystallogr 66, 486–501 (2010).

61. Kelley, L.A., Mezulis, S., Yates, C.M., Wass, M.N. & Sternberg, M.J. The Phyre2 web portal for protein modeling, prediction and analysis. Nat Protoc 10, 845–58 (2015).

62. Goddard, T.D. et al. UCSF ChimeraX: Meeting modern challenges in visualization and analysis. Protein Sci 27, 14–25 (2018).

63. Lindsley, J.E. & Wang, J.C. On the coupling between ATP usage and DNA transport by yeast DNA topoisomerase II. J Biol Chem 268, 8096–104 (1993).

