## Supplementary Information for "Cryo-EM structure of the complete *E. coli* DNA Gyrase nucleoprotein complex"

### TABLE OF CONTENT


**pg. 2** Figure S1. Purification and DNA relaxation activity of reconstituted *E. coli* DNA Gyrase

**pg. 3** Figure S2. Cryo-EM data acquisition and *ab-initio* model generation

**pg. 4-5** Figure S3. Flow chart of the Cryo-EM data processing

**pg. 6** Figure S4. Cryo-EM statistics of the overall DNA-bound *E. coli* DNA Gyrase and DNA-binding and cleavage core in closed and pre-opening states

**pg. 7** Figure S5. High and intermediate resolution cryo-EM maps of the different functional domains fitted into the low resolution cryo-EM map of the overall complex

**pg. 8** Figure S6. Comparison of the X-ray and cryo-EM structures of *E. coli* DNA Gyrase at the GyrB-GyrA extremities interface

**pg. 9** Figure S7. DNA Gyrase overall geometry analysis

**pg. 10** Figure S8. Conservation of the GHKL/Transducer domains and R286 residue in bacteria

**pg. 11** Figure S9. Thermal stability analysis and DNA stimulated ATP hydrolysis activity

**pg. 12-13** Figure S10. Quaternary and tertiary changes associated with G-segment binding and opening after cleavage

**pg. 14** Figure S11. Metal-binding site associated with cleavage of DNA by DNA Gyrase

**pg. 15** Figure S12. CTD β-pinwheel acidic tail modeling

**pg. 16** Figure S13. Comparison of the *E. coli* and *S. aureus* NBTI binding site

**pg. 17** Figure S14. Experimental map quality around the NBTI molecule

**pg. 18** Table S1. Asymmetric oligonucleotides sequences.

**pg. 19** Table S2. Data collection, processing and refinement statistics

**pg. 20** Table S3. Missing DNA gyrase sequence elements modeled in this study

**pg. 21** Supplementary references

### SUPPLEMENTARY FIGURES

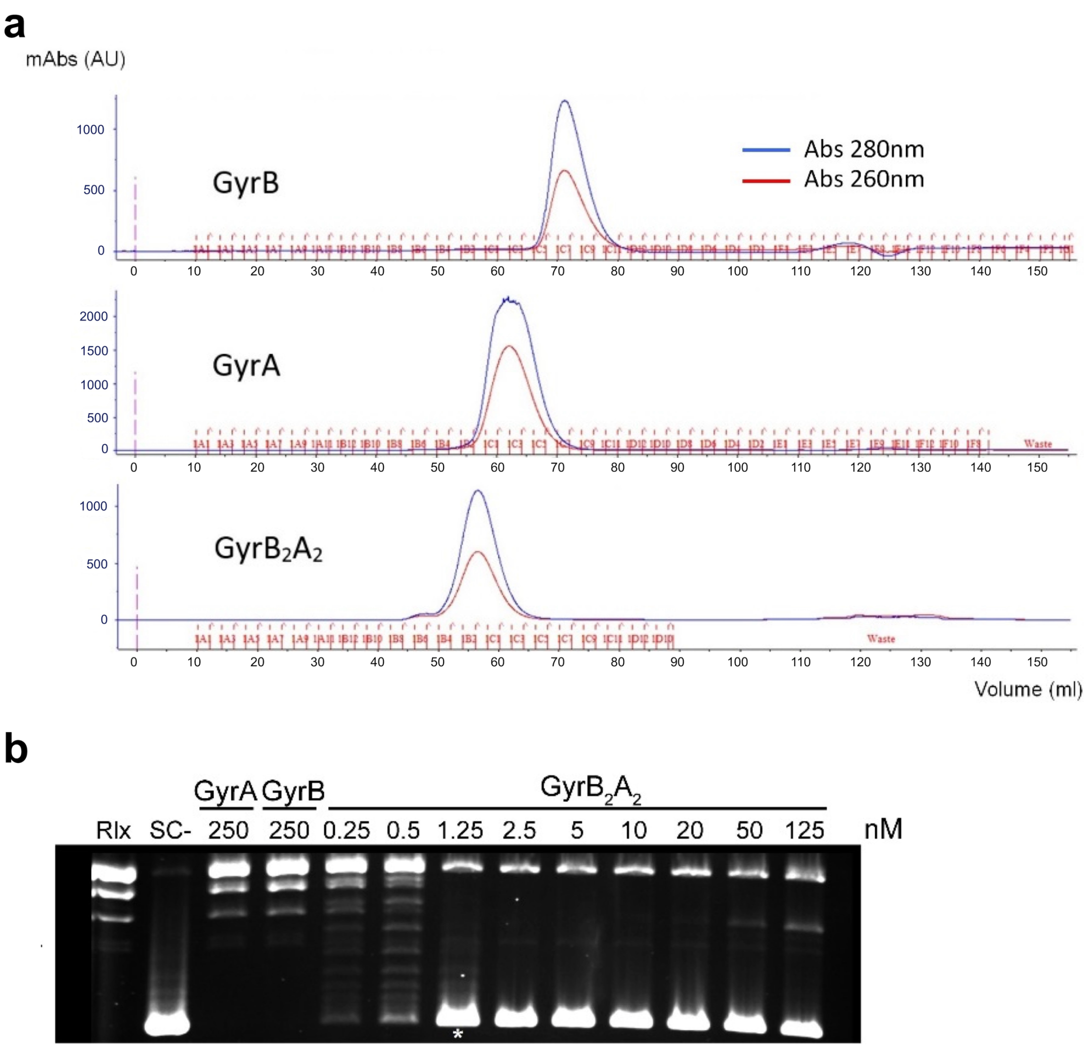


**Figure S1. Purification of reconstituted *E. coli* DNA Gyrase. a.** Comparison of the elution volumes of the GyrB subunit, GyrA subunit and the reconstituted $GyrA_2B_2$ DNA Gyrase showing a clear shift on a gel filtration column (Superdex S200 16/60). **b.** Negative supercoiling activity of the reconstituted *E. coli* DNA Gyrase. Protein concentrations are listed in nM holoenzyme and asterisk indicates concentration of enzyme needed to supercoil the substrate in 30 min. Negative and positive controls are shown as relaxed (Rlx) or negatively supercoiled DNA species (SC-), respectively.

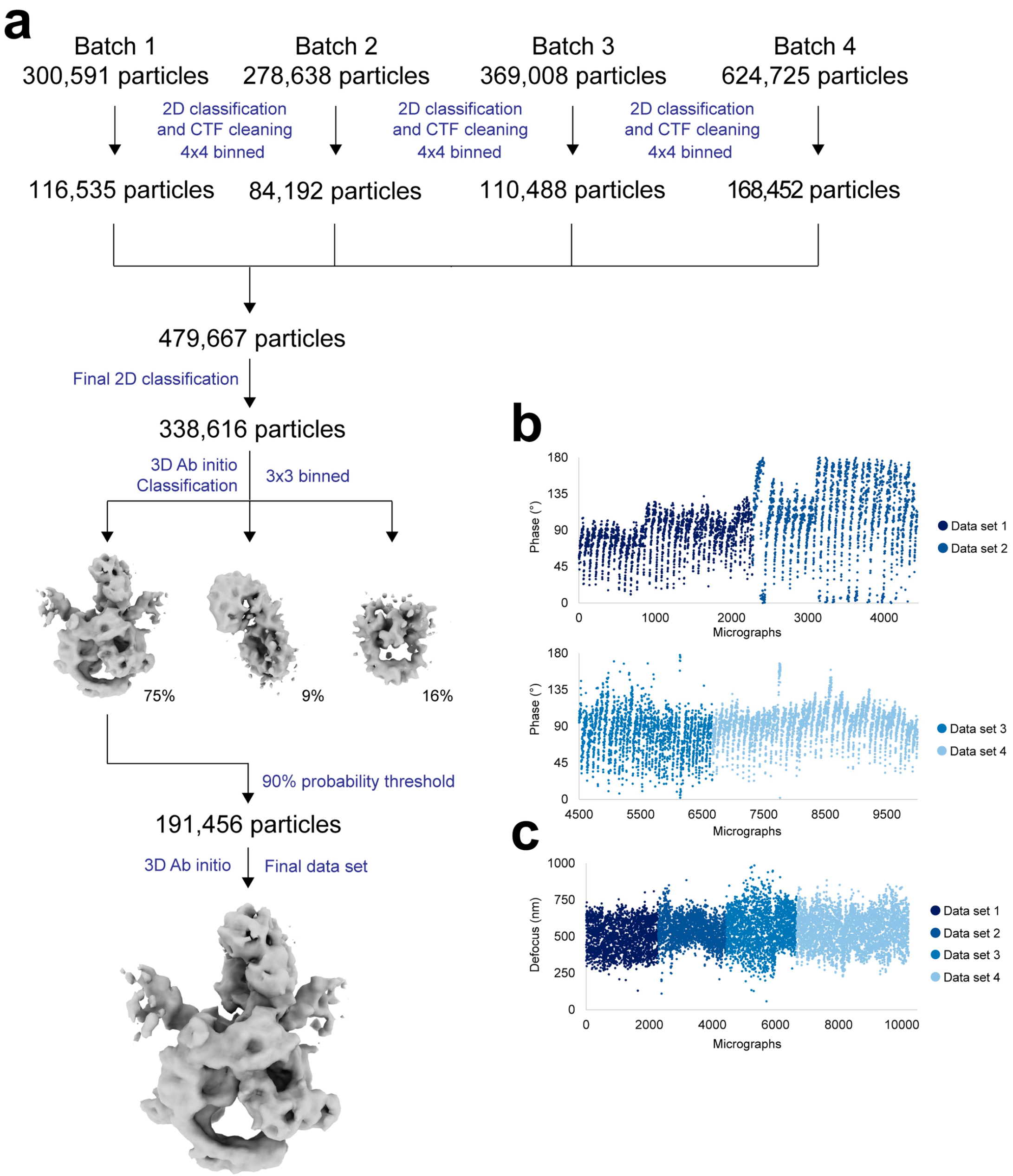


**Figure S2. Cryo-EM data acquisition and *ab-initio* model generation. a.** Flow chart of data processing from 2D classification to *ab-initio* model generation. The particle numbers are indicated at each step. **b.** Volta phase plate phase shift throughout the dataset. **c.** Defocus estimation history throughout the dataset.

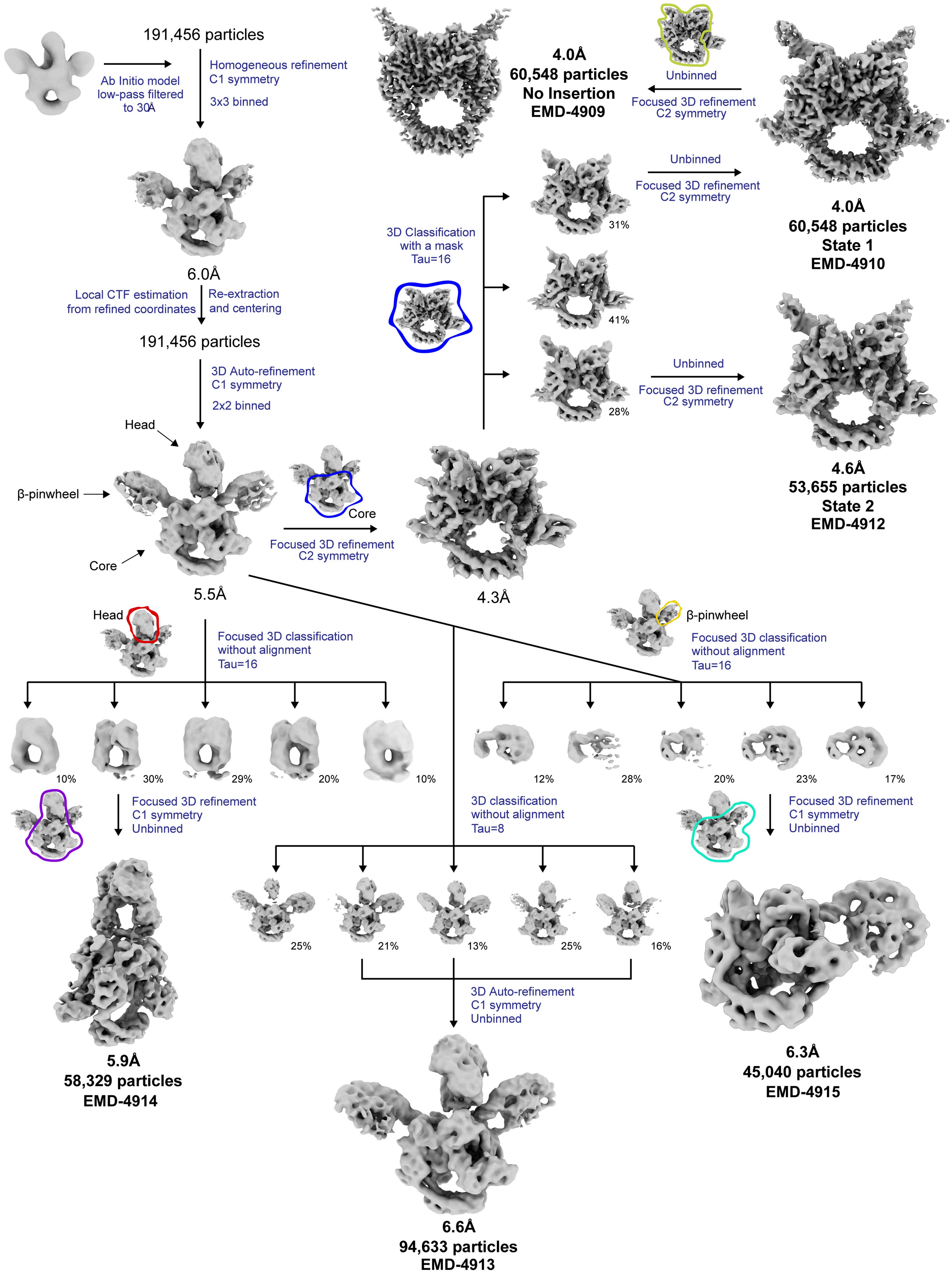


**Figure S3. Flow chart of the Cryo-EM data processing.** *Ab-initio* model was refined in cryoSPARC [1] to 6.0 Å overall using 191,456 particles. Using the refined coordinates, particles were re-extracted and centered and per-particle CTF estimation was performed with GCTF [2]. Using this new particles stack, the previous structure was refined to 5.5 Å overall. Using RELION2 [3,4], focused 3D classification with and without alignment followed by focused 3D refinement allowed to solve 6 new structures: the DNA-binding and cleavage domain in closed state (with and without TOPRIM insertion) at 4.0 Å and pre-opening state at 4.6 Å, the DNA-binding and cleavage domain with the ATPase domain at 5.9 Å, the DNA-binding and cleavage domain with the β-pinwheel 6.3 Å and the overall complex with improved quality of the flexible regions at 6.6 Å. The particle numbers for the final refinement are indicated. 3x3, 2x2 binned and unbinned structures corresponds to pixel sizes of 2.64, 1.76 and 0.88 Å/px, respectively.

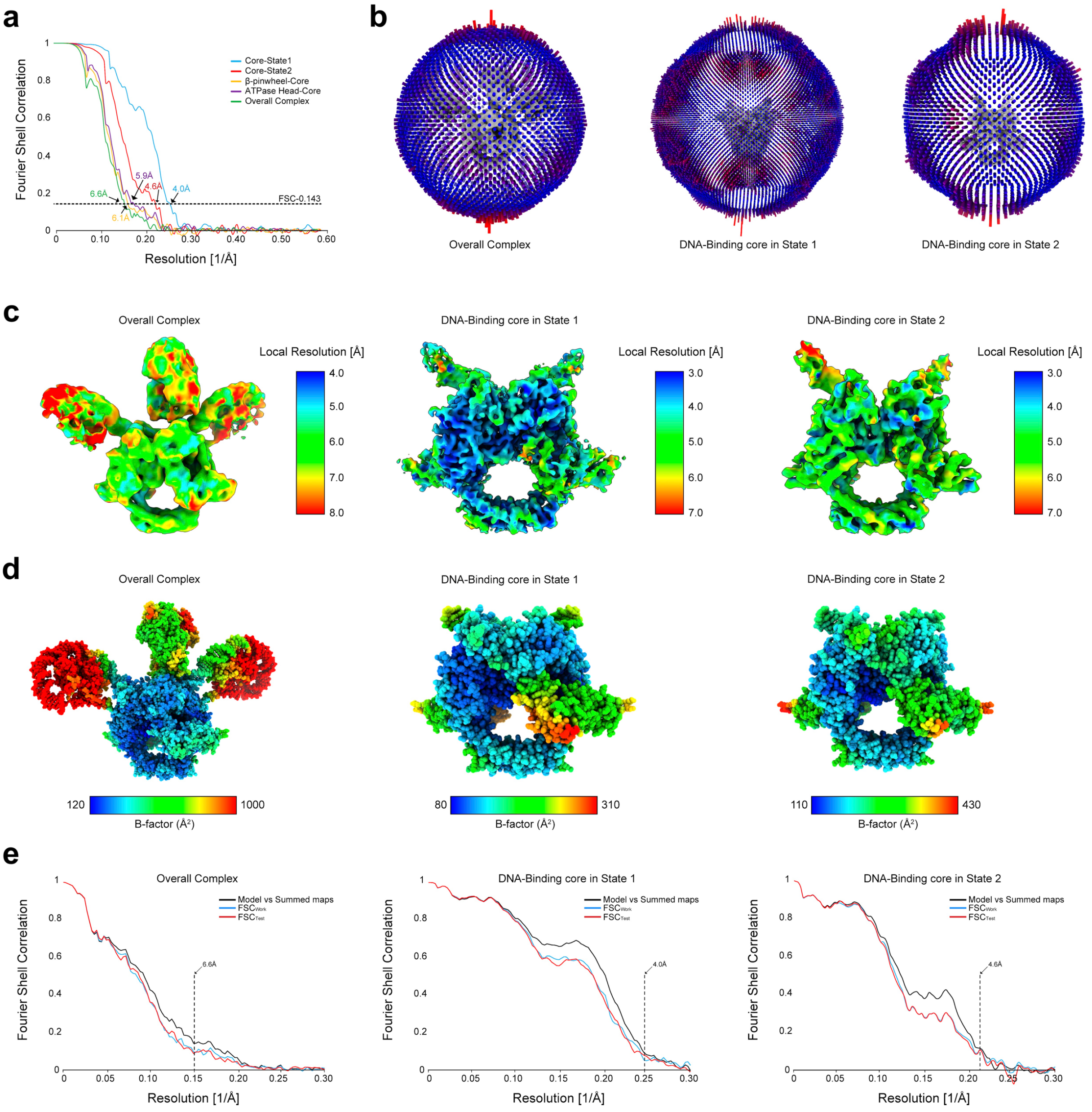


**Figure S4. Cryo-EM statistics of the overall DNA-bound *E. coli* DNA Gyrase and DNA-binding and cleavage core in closed and pre-opening states. a.** FSC plots and resolution estimation using the gold-standard 0.143 criterion generated from RELION2 [3,4]. **b.** Angular distribution plots generated from RELION2. **c.** Final refined map colored according to local resolution calculated with Blocres [5]. **d.** Atomic models refined in the corresponding cryo-EM maps colored according to the B-factors. **e.** Cross-validation FSC curves for the corresponding refined models versus unfiltered half maps (the one used in the refinement, $FSC_{work}$, and the other half, $FSC_{free}$) and the unfiltered summed maps.

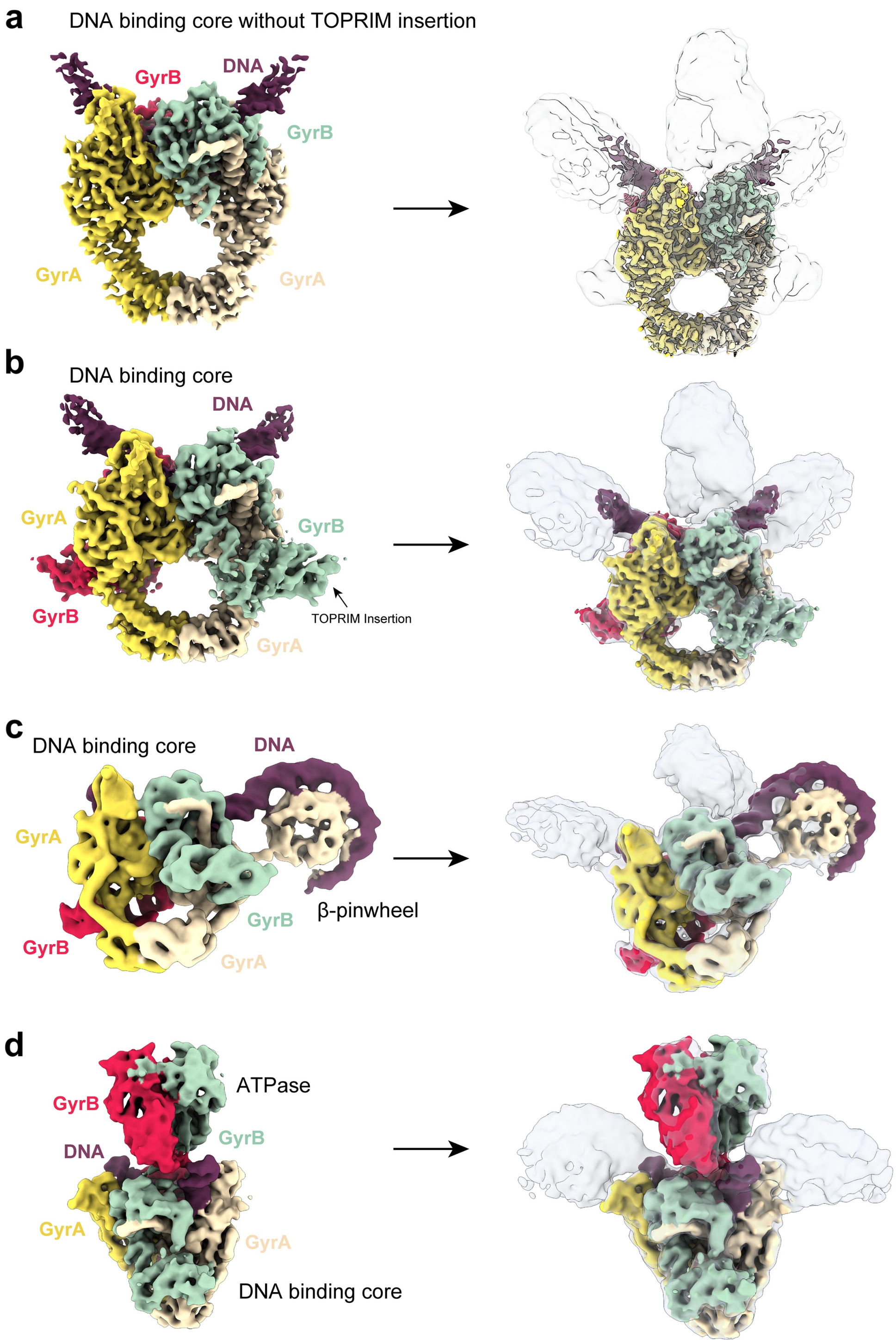


**Figure S5. High and intermediate resolution cryo-EM maps of the different functional domains fitted into the low resolution cryo-EM map of the overall complex. a.** The DNA-binding and cleavage domain in closed state lacking the TOPRIM insertion solved at 4 Å resolution fitted in the 6.6 Å map of the overall complex. **b.** The DNA-binding and cleavage domain in closed state solved at 4 Å resolution fitted in the 6.6 Å map of the overall complex. **c.** The DNA-binding and cleavage domain in closed state with the β-pinwheel domain solved at 6.3 Å resolution fitted in the 6.6 Å map of the overall complex. **d.** The DNA-binding and cleavage core in closed state with the ATPase domain solved at 5.9 Å resolution fitted in the 6.6 Å map of the overall complex. The different maps were used to build and refine the atomic model of the entire complex.

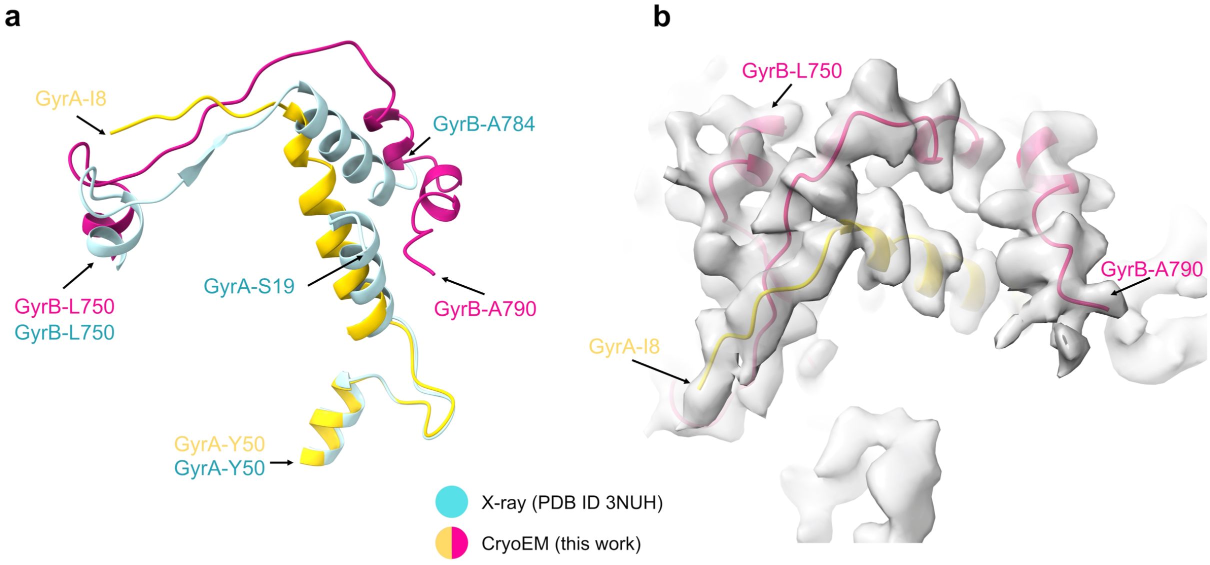


**Figure S6. Comparison of the X-ray and cryo-EM structures of *E. coli* DNA Gyrase at the GyrB-GyrA extremities interface. a.** After several rounds of manual building and refinement, the resulting cryo-EM atomic model was superimposed on the X-ray structure (PDB ID 3NUH) [6] in the area of the GyrB C-terminal (L750-A790) and GyrA N-terminal (I8-Y50) extremities. The superimposition of the structures shows a misalignment of both GyrB and GyrA extremities. **b.** The quality of the cryo-EM map allowed to unambiguously build and correct this area.

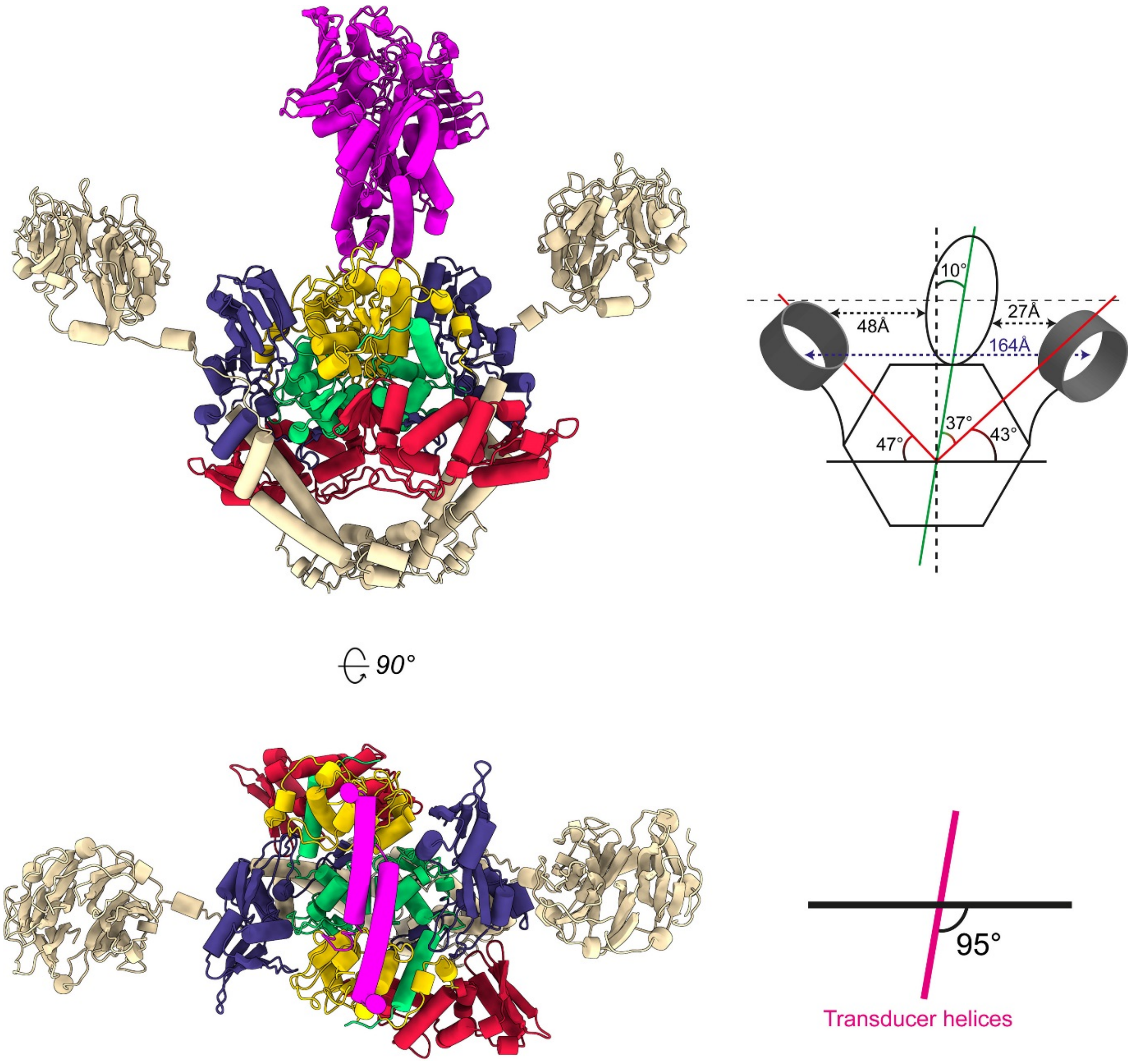


**Figure S7. DNA Gyrase overall geometry analysis.** The domains are colored as follow: ATPase in magenta, TOPRIM in yellow, TOPRIM insertion in red, WHD in green, Tower in blue, Coil-coiled and CTD in beige. DNA is omitted for clarity. Upper panel: a schematic representation of the domains' orientation is shown for the gyrase complex model. The DNA-binding and cleavage domain (DNA- and C-gate) is depicted as a hexagon, the GyrB ATPase domain (N-gate) as an ellipse, and the CTD β-pinwheel domains as disks. Red and green solid lines indicate the β-pinwheel and ATPase domain planes, respectively. The ATPase domain bends toward one β-pinwheel with an angle of ~10°. This positions the ATPase domain at a distance of 26 Å from the β-pinwheel. The second β-pinwheel is 48Å from ATPase domain. The β-pinwheels are located in an upper position and are distributed asymmetrically on each side of the DNA gate (47° versus 43°). Lower panel: Top view of the structure with the upper part of the ATPase domain omitted. The ATPase domain transducer α-helices in magenta and forms a ~95° angle with the DNA binding-cleavage domain.

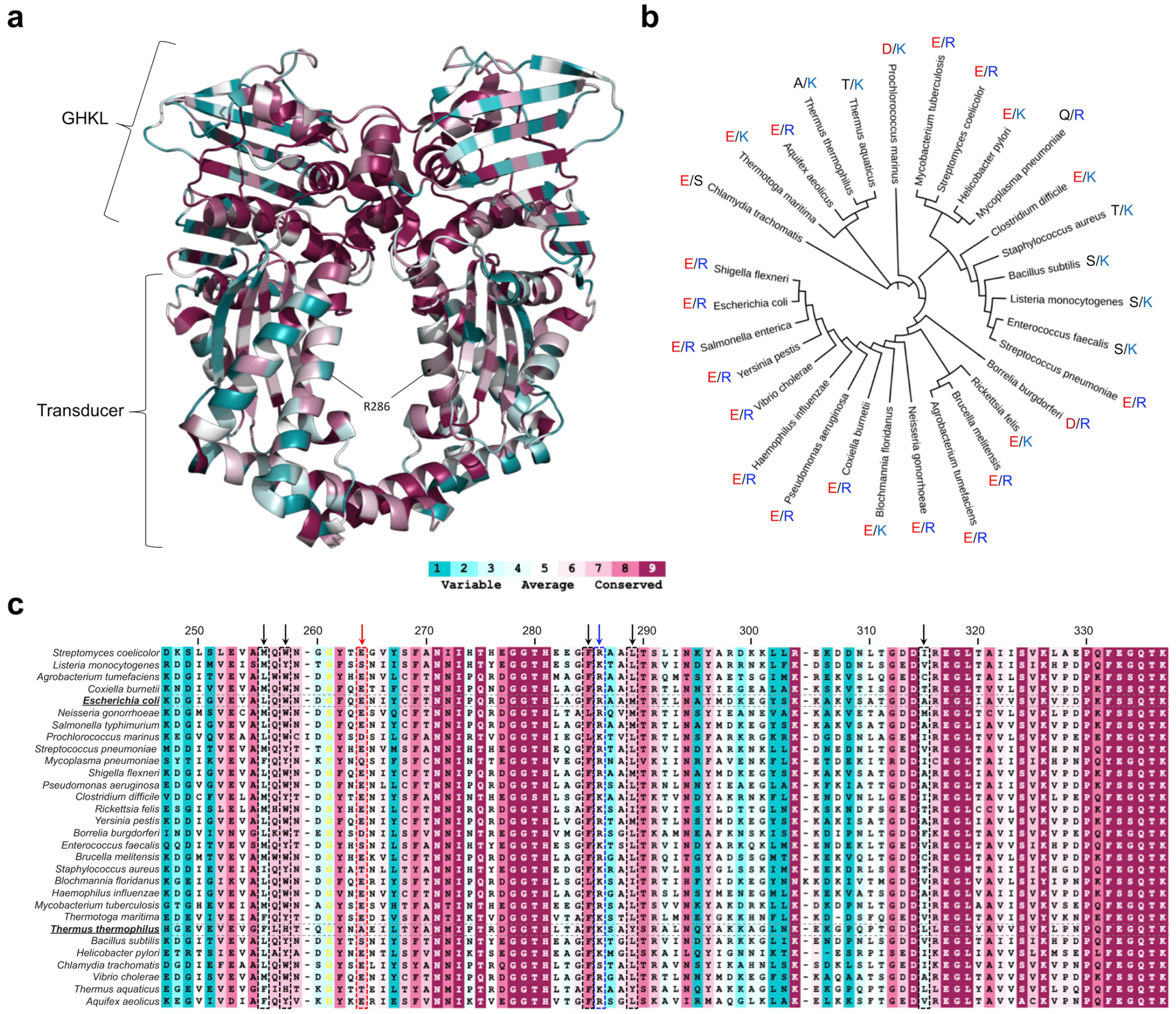


**Figure S8. Conservation of the GHKL/Transducer domains and R286 residue in bacteria. a.** Atomic model of the ATPase and transducer domains (PDB ID 1EI1) [7] colored by conservation level for each residue among 30 species within Gram-positive and Gram-negative bacteria. All the residues interacting with ATP in the GHKL domain are highly conserved while residues in the transducer show a smaller degree of conservation. **b.** Phylogenetic tree generated using a multiple alignment of the ATPase/transducer sequences from 30 bacterial species (listed in c). The R286-E264 couple (numbering based on *E. coli* sequence) is indicated for each species. R286 is found either as an Arginine, mostly in Gram negative bacteria, or as a Lysine in Gram positive bacteria. **c.** Multiple alignment of the ATPase/Transducer domain focused on the transducer domain. Black arrows indicate the residues implicated in hydrophobic contacts stabilizing the transducer beta-strands and alpha helices. The blue arrow indicates the R286 and homologous residues within the 30 species. The red arrow indicates the E264 and homologous residues within the 30 species.

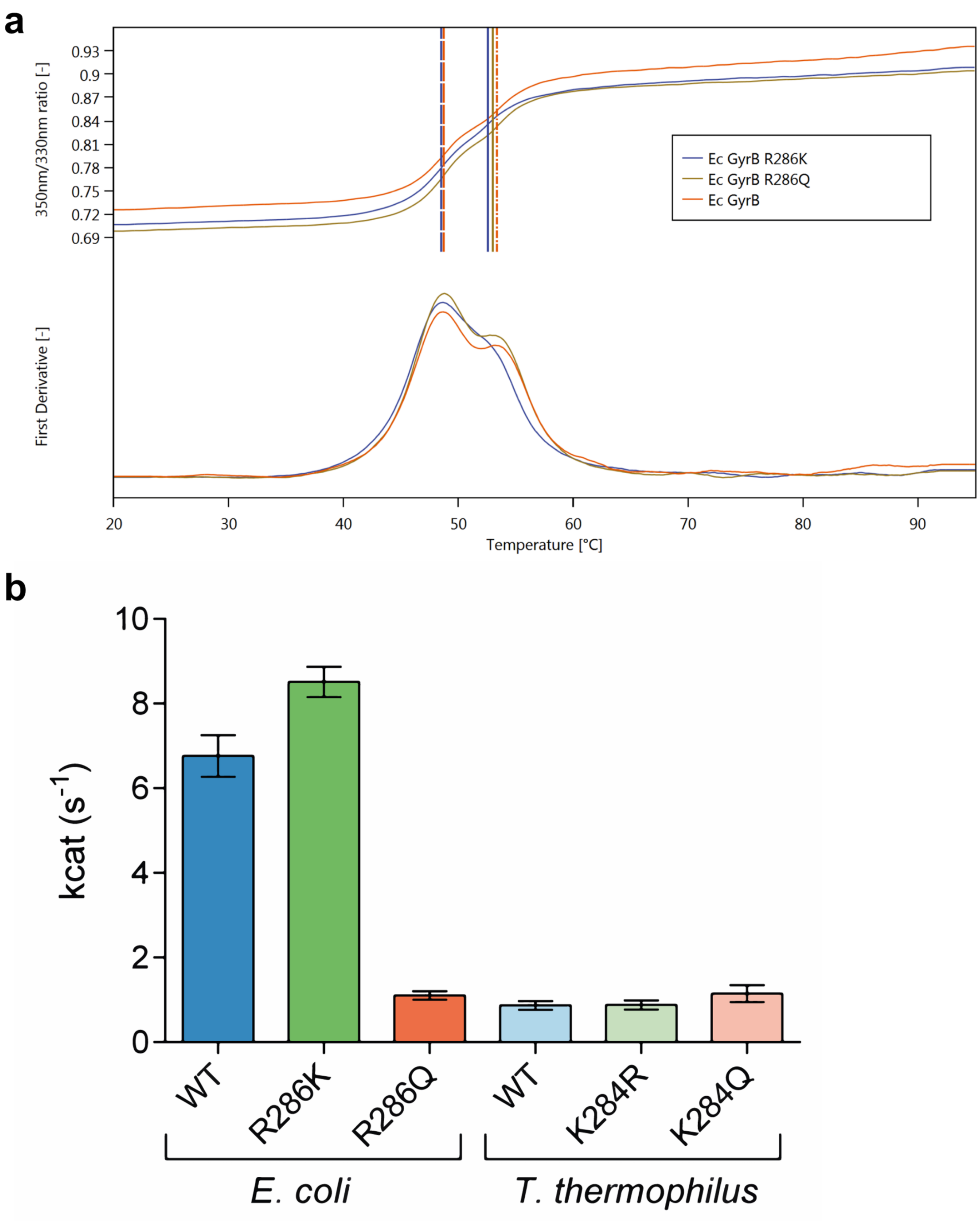


**Figure S9. Thermal stability analysis and DNA stimulated ATP hydrolysis activity**. **a**. The thermal stability of the *E. coli* WT, R286K and R286Q GyrB domains was measured by differential scanning fluorimetry based on the intrinsic fluorescence of tryptophan. The denaturation curve of the WT GyrB and of the mutants shows no major difference (upper panel). The first derivative (lower panel) displays two peaks consistent with a multidomain protein with no significant shift of the main temperature transition peak. **b.** DNA stimulated ATP hydrolysis activity of WT, R286K and R286Q *E. coli* DNA gyrase and WT, K284R and K284Q *T. thermophilus* DNA gyrase. The mutation R286K shows no negative effect on the ATPase activity of the *E. coli* DNA Gyrase. The R286Q shows a 7-fold reduction of its ATPase activity. The K284R and K284Q mutations have no effect on ATPase activity of *T. thermophilus* DNA Gyrase.

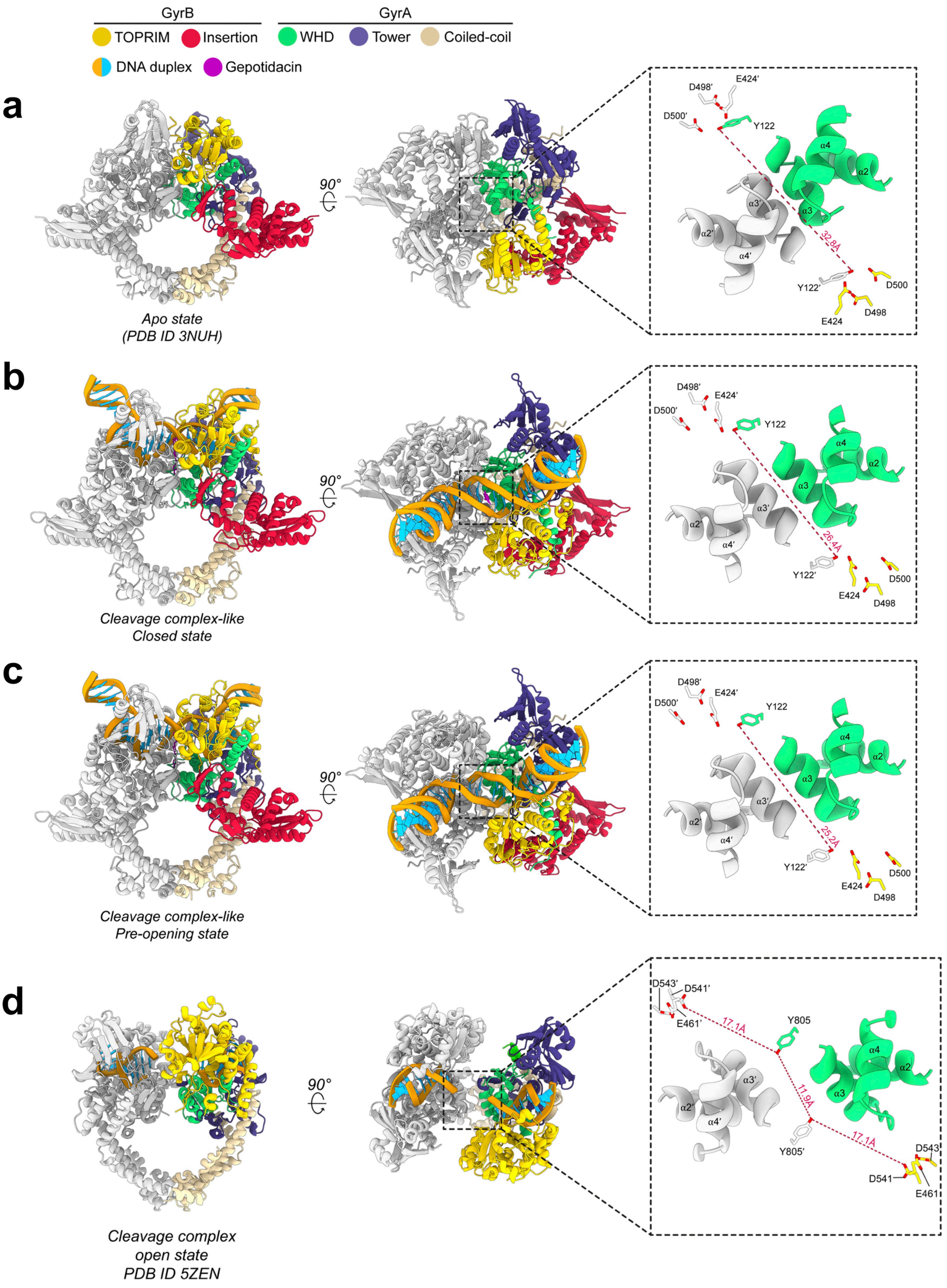


**Figure S10. Quaternary and tertiary changes associated with G-segment binding and opening after cleavage.** Orthogonal views of the *E. coli* DNA-binding and cleavage core in different states. The inset shows selected structural elements (alpha helices α2, α3 and α4) lining the bottom side of the G-segment groove and key catalytic groups in the *apo* [6] (A) CC closed (B) and CC pre-opening (C) and open [8] (D) conformations. For each conformation, the catalytic tyrosines, the $Mg^{2+}$-binding residues, and the distance between the two catalytic tyrosines are shown to illustrate structural changes in the DNA-gate during the Apo-to-closed and closed-to-pre-opening transitions. Residues from different homodimer (GyrB-GyrA or hsTop2β) are colored differently and marked by a quotation mark.

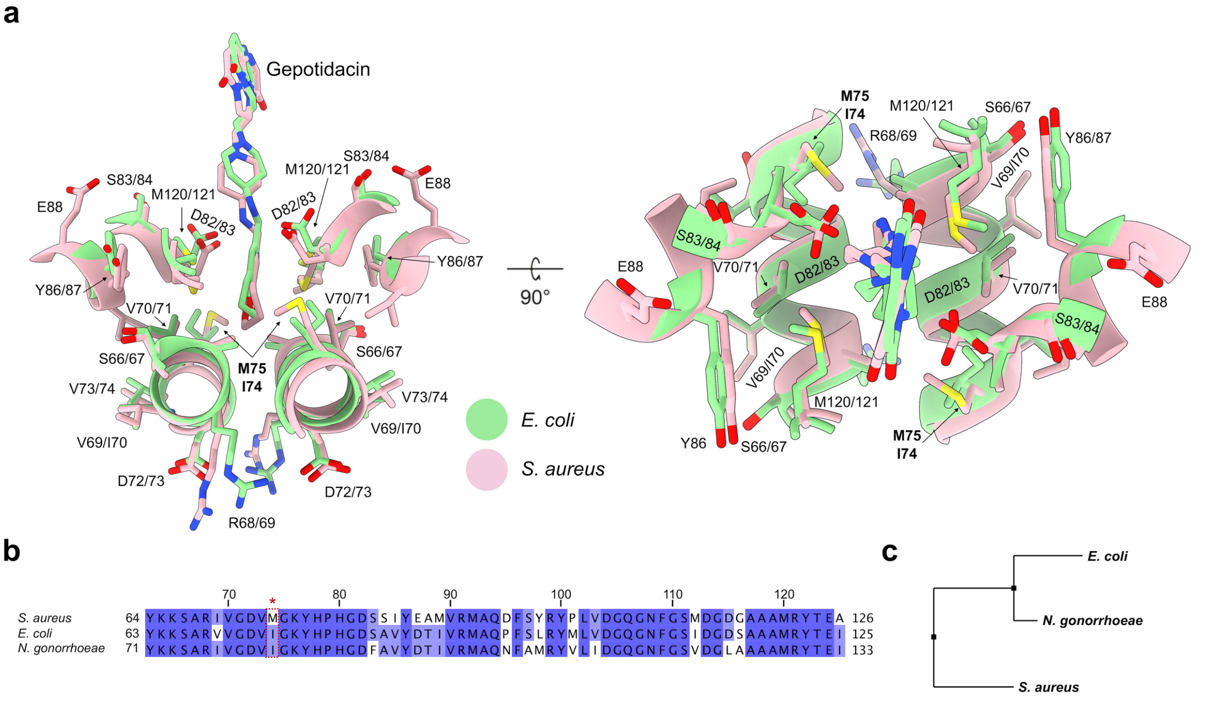


**Figure S11. Comparison of the *E. coli* and *S. aureus* NBTI binding site. a.** Superimposition of the NBTI binding sites from the *E. coli* closed conformation (this study) in green and from the crystal structure of the *S. aureus* DNA binding and cleavage domain in pink (PDB ID 6QTK) [9]. Residues in the close vicinity of the NBTI are annotated. The first number corresponds to the *E. coli* residue and the second to the *S. aureus* residue. The major difference resides on the I74 in *E. coli* that corresponds to a Methionine residue (Met75) in *S aureus.* **b.** Sequence alignment on the NBTI binding region of *E. coli* (Uniprot: P0AES6), *S. Aureus* (Uniprot: P0A0K8) and *N. Gonorrhoeae* (Uniprot: P22118). **c.** Phylogenetic tree generated using the multiple alignment from (b). *N. Gonorrhoeae* and *E. coli* are evolutionary closer than *S. aureus*.

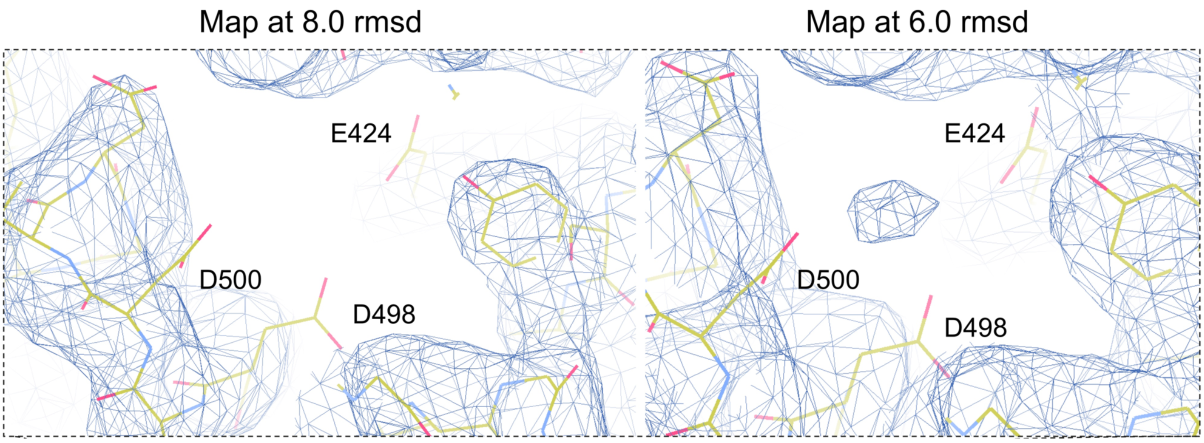


**Figure S12. Metal-binding site associated with cleavage of DNA by DNA Gyrase.** EM density of the cleavage-complex structure solved at 4 Å resolution at different rsmd steps of contouring showing a density at the site of a $Mg^{2+}$ ion at 6.0 rmsd.

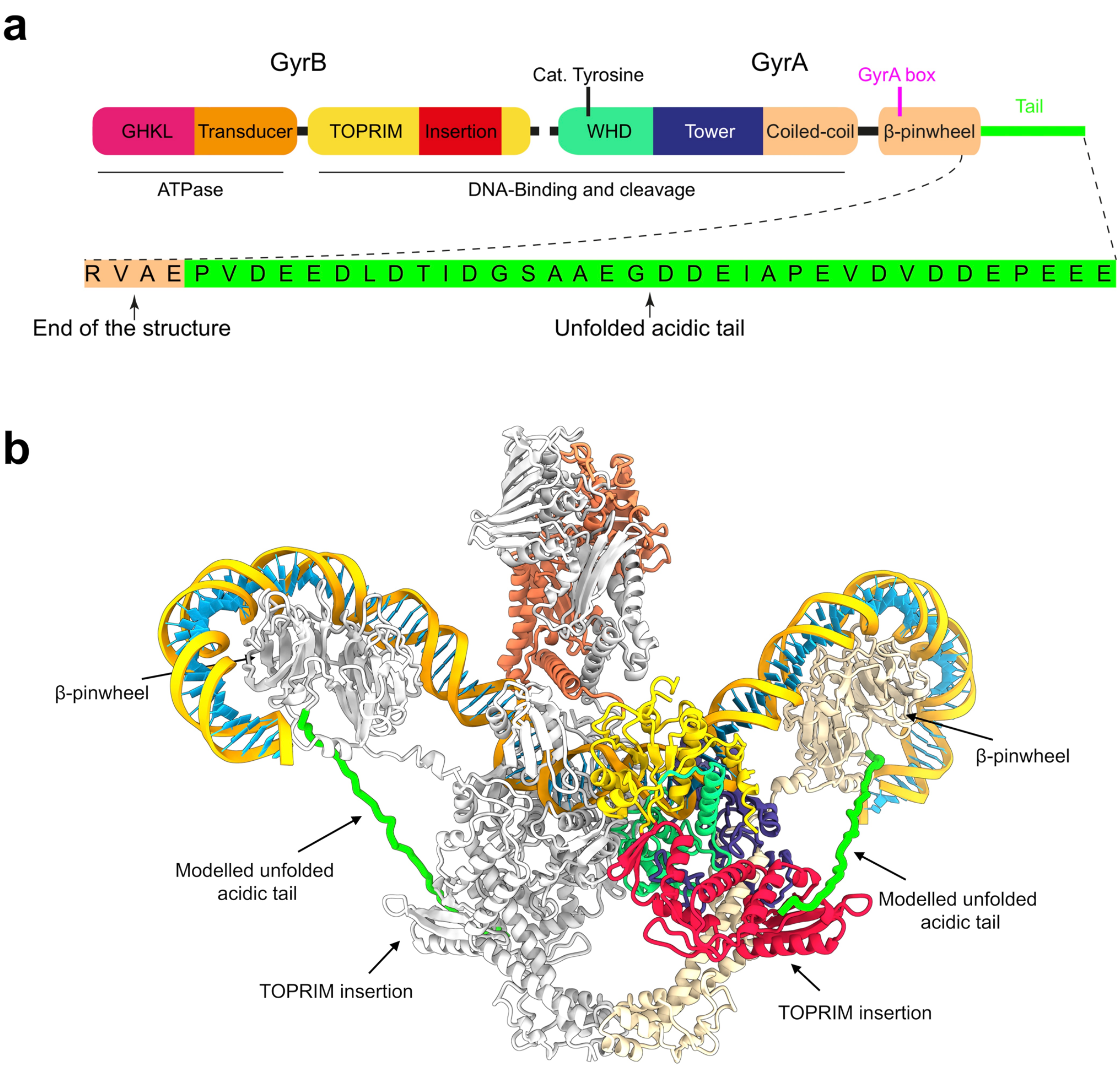


**Figure S13. CTD β-pinwheel acidic tail modeling. a.** Primary domain structure of the *E. coli* DNA Gyrase. The β-pinwheel acidic tail is highlighted in green. Sequence of the *E. coli* GyrA β-pinwheel tail is shown in the corresponding colors. **b.** Structure of the *E. coli* DNA Gyrase complex with the 130bp DNA duplex. The unfolded β-pinwheel acidic tails were modelled as coils (green). The distance between the C-terminal end of the β-pinwheel and the GyrB TOPRIM insertion domain is compatible with the 34 residues sequence of the tail, triggering potential contact between the 2 domains.

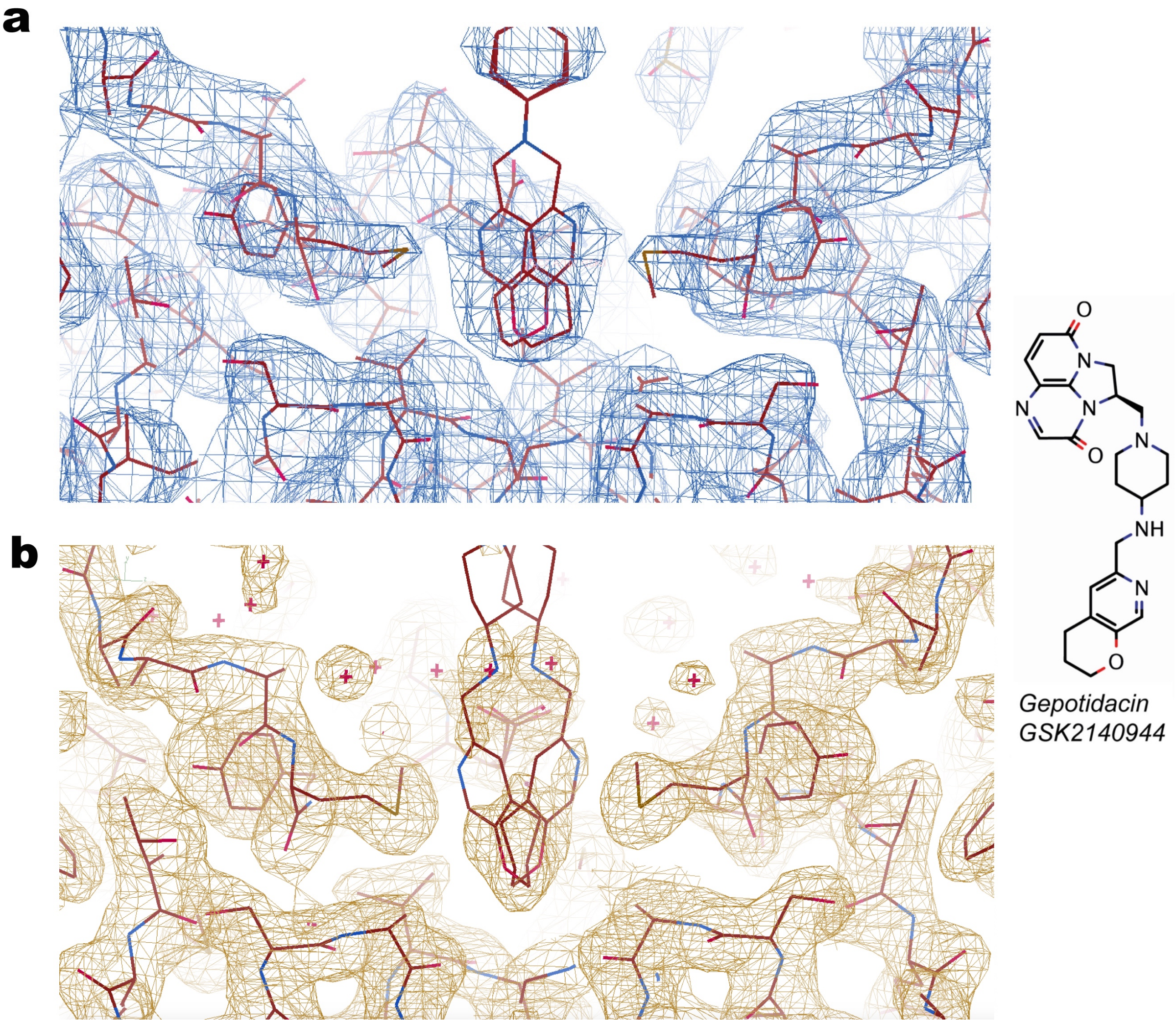


**Figure S14. Experimental map quality around the NBTI molecule. a.** Cryo-EM map at 1.6 rmsd of the 4 Å structure of the DNA binding and cleavage domain in closed state (our study). **b.** Electron density map contoured at 1 σ from the crystal structure of the *S. aureus* DNA binding and cleavage domain (PDB ID 6QTK) [9]. The same level of details can be obtained in this region when comparing the X-ray and the cryo-EM data.

### SUPPLEMENTARY TABLES

| | |
|---|---|
| 92bp-AsymDuplex | GTACCTACGGCTTATGATTCTTCTCGCTTCCGGCGGCATCGG GATGCCCGCGTTGCAGGCCATGCTGTCCAGGCAGGTAGATGA CGACCATC |
| 88bp-AsymDuplex | GATGGTCGTCATCTACCTGCCTGGACAGCATGGCCTGCAACG CGGGCATCCCGATGCCGCCGGAAGCGAGAAGAATCATAAGC CGTAG |

**Table S1. Asymmetric oligonucleotides sequences.**

| **Data Collection** | |
|---|---|
| Microscope | Titan Krios |
| Voltage (keV) | 300 |
| Magnification | 130,000 |
| Electron dose ($e^{-}$ Å$^{-2}$) | 50 |
| Detector | Gatan K2 Summit (super-resolution mode) |
| Pixel Size (Å) | 0.88 (0.44) |
| Defocus target (nm) | -500 |

| **Individual data sets** | | | | |
|---|---|---|---|---|
| Batches | **1** | **2** | **3** | **4** |
| Micrographs (no.) | 3475 | 2528 | 2922 | 3980 |
| Extracted particles (no.) | 300,591 | 278,638 | 369,008 | 624,725 |
| Particles after cleaning | 116,535 | 84,192 | 110,488 | 168,452 |

| **Merged data sets for refinement** | |
|---|---|
| Total merged particles | 479,667 |
| Particles after cleaning (one round of 2D and 3D ab initio classification) | 192,456 |

| **Reconstruction** | | | | | | |
|---|---|---|---|---|---|---|
| | **Closed Core ΔInsertion** | **Closed Core** | **Pre-opening Core** | **Overall complex** | **ATPase & Core** | **CTD & Core** |
| EMDB | EMD-4909 | EMD-4910 | EMD-4912 | EMD-4913 | EMD-4914 | EMD-4915 |
| PDB | 6RKS | 6RKU | 6RKV | 6RKW | - | - |
| Software | Relion2.0.3 | | | | | |
| Final particles (no.) | 60,548 | 60,548 | 53,655 | 94,633 | 58,329 | 45,040 |
| Box size (pixels) | 360x360x360 | | | | | |
| Symmetry imposed | C2 | C2 | C2 | C1 | C1 | C1 |
| Map resolution FSC 0.143 (global) (Å) | 4.0 | 4.0 | 4.6 | 6.6 | 5.9 | 6.3 |
| Applied B-factor for sharpening (Å$^2$) | -144.9 | -79.4 | -167.2 | -246.5 | -234.6 | -144.9 |

| **Model refinement** | | | | | | |
|---|---|---|---|---|---|---|
| Software | Phenix 1.12-2829 | | | | - | - |
| Resolution cut-off (Å) | 4.0 | 4.0 | 4.6 | 6.6 | | |
| Unit cell (Å) | 316.8x316.8x316.8 | | | | | |
| Non-hydrogen atoms | 12572 | 15526 | 15460 | 30270 | | |
| Protein residues | 1416 | 1784 | 1784 | 3218 | | |
| DNA bases (atoms) | 1312 | 1312 | 1312 | 4920 | | |
| Ligands (atoms) | 66 | 66 | 0 | 128 | | |
| Average B-factor | 69.03 | 145.1 | 187.4 | 532.8 | | |
| R.m.s. deviations | | | | | | |
| Bond lengths (Å) | 0.004 | 0.004 | 0.003 | 0.003 | - | - |
| Bond angles (°) | 0.936 | 0.904 | 0.877 | 0.839 | | |

| **Validation** | | | | | | |
|---|---|---|---|---|---|---|
| Real space correlation coefficient (Global) | 0.83 | 0.84 | 0.80 | 0.77 | - | - |
| MolProbity score | 1.78 | 1.80 | 1.76 | 1.61 | | |
| Clashscore (all atoms) | 5.24 | 5.93 | 7.23 | 7.62 | | |
| Poor rotamers (%) | 0 | 0 | 0 | 0.07 | | |
| Ramachandran plot | | | | | | |
| Favoured (%) | 91.64 | 92.35 | 94.67 | 96.83 | - | - |
| Allowed (%) | 8.36 | 7.65 | 5.33 | 3.17 | | |
| Outliers (%) | 0 | 0 | 0 | 0 | | |

**Table S2. Data collection, processing and refinement statistics**.

| Uniprot ID | Protein name | Domain | PDB ID | Residues range | Comment | Reference |
|---|---|---|---|---|---|---|
| GYRB_ECOLI | GyrB Total of 37 residues added | ATPase | 1EI1 | 2-392 | | [7] |
| | | Linker | | 393-401 (9) | manually built | this work |
| | | DNA binding and cleavage | 3NUH | 402-448 | | [6] |
| | | | | 449-461 (13) | manually built | this work |
| | | | 3NUH | 462-475 | | [6] |
| | | | | 483-488 (6) | manually built | this work |
| | | | 3NUH | 489-655 | | [6] |
| | | | | 656-658 (3) | manually built | this work |
| | | | 3NUH | 659-765 | | [6] |
| | | | | 766-784 | 3NUH manually corrected (Figure S6) | this work |
| | | | | 785-790 (6) | manually built | this work |
| GYRA_ECOLI | GyrA Total of 49 residues added | DNA binding and cleavage | | 8-18 (11) | manually built | this work |
| | | | 3NUH | 19-174 | | [6] |
| | | | | 175-178 (4) | manually built | this work |
| | | | 3NUH | 179-251 | | [6] |
| | | | | 252-255 (4) | manually built | this work |
| | | | 3NUH | 256-415 | | [6] |
| | | | | 416-419 (4) | manually built | this work |
| | | | 3NUH | 420-422 | | [6] |
| | | | | 427-429 (3) | manually built | this work |
| | | | 3NUH | 430-441 | | [6] |
| | | | | 442-443+446 (3) | manually built | this work |
| | | | 3NUH | 447-524 | | [6] |
| | | Linker | | 525-534 (10) | manually built | this work |
| | | β-pinwheel | 1ZI0 (chain B) | 535-563 | | [10] |
| | | | | 564-574 (11) | modeled by Phyre2 | this work |
| | | | 1ZI0 (chain B) | 575-841 | | [10] |

**Table S3. Missing DNA gyrase sequence elements modeled in this study.**

### Supplementary References


1. Punjani, A., Rubinstein, J.L., Fleet, D.J. & Brubaker, M.A. cryoSPARC: algorithms for rapid unsupervised cryo-EM structure determination. *Nat Methods* **14**, 290-296 (2017).
2. Zhang, K. Gctf: Real-time CTF determination and correction. *J Struct Biol* **193**, 1-12 (2016).
3. Scheres, S.H. RELION: implementation of a Bayesian approach to cryo-EM structure determination. *J Struct Biol* **180**, 519-30 (2012).
4. Kimanius, D., Forsberg, B.O., Scheres, S.H. & Lindahl, E. Accelerated cryo-EM structure determination with parallelisation using GPUs in RELION-2. *Elife* **5**(2016).
5. Heymann, J.B. & Belnap, D.M. Bsoft: image processing and molecular modeling for electron microscopy. *J Struct Biol* **157**, 3-18 (2007).
6. Schoeffler, A.J., May, A.P. & Berger, J.M. A domain insertion in Escherichia coli GyrB adopts a novel fold that plays a critical role in gyrase function. *Nucleic Acids Res* **38**, 7830-44 (2010).
7. Brino, L. et al. Dimerization of Escherichia coli DNA-gyrase B provides a structural mechanism for activating the ATPase catalytic center. *J Biol Chem* **275**, 9468-75 (2000).
8. Chen, S.F. et al. Structural insights into the gating of DNA passage by the topoisomerase II DNA-gate. *Nat Commun* **9**, 3085 (2018).
9. Gibson, E.G., Bax, B., Chan, P.F. & Osheroff, N. Mechanistic and Structural Basis for the Actions of the Antibacterial Gepotidacin against Staphylococcus aureus Gyrase. *ACS Infect Dis* **5**, 570-581 (2019).
10. Ruthenburg, A.J., Graybosch, D.M., Huetsch, J.C. & Verdine, G.L. A superhelical spiral in the Escherichia coli DNA gyrase A C-terminal domain imparts unidirectional supercoiling bias. *J Biol Chem* **280**, 26177-84 (2005).
